# A portable C-terminal rheostat calibrates RecE/Redα exonuclease activity for enhanced bacterial recombineering

**DOI:** 10.64898/2026.10.29.762211

**Authors:** Ji Luan, Chanjuan Jiang, Ruofei Guo, Chongyang Wang, Shaoqian Wang, A. Francis Stewart, Jun Fu, Youming Zhang, Hailong Wang

**Author notes:** J.L., C.J., R.G., and C.W. contributed equally to this work. Correspondence should be addressed to Hailong Wang.

## Abstract

Recombineering enables precise genome engineering across diverse bacteria, yet efficient tools for large DNA integration remain underdeveloped for Streptomyces and other biotechnologically relevant bacteria. Here we identify a panel of highly active RecET recombinase pairs from actinobacteriophages that mediate efficient homologous recombination in Streptomyces. The lead candidate, Che9cRecET, supports genomic integration of kilobase-scale DNA with only 50bp homology arms and enables accurate insertion of 79kb biosynthetic gene clusters, far outperforming conventional genome editing approaches. Structural and biochemical analyses reveal Che9cRecE as a distinct C3-symmetric toroidal homotrimeric 5′-3′ exonuclease, whose extended C-terminal loop acts as a tunable rheostat calibrating exonuclease activity to an optimal window for recombination. We further show this portable C-terminal loop enhances recombineering efficiency across phylogenetically diverse exonucleases including Escherichia coli Redα/RecE and Pseudomonas A. Our work uncovers a conserved regulatory mechanism of the RecE/Redα exonuclease family and delivers a generalizable strategy for advancing bacterial genome editing.

## Introduction

Recombineering is a broadly applicable bacterial genome-editing technology mediated by 5′–3′ double-stranded DNA (dsDNA) exonucleases (Redα or RecE) and their cognate single-stranded DNA (ssDNA) annealing proteins (SSAPs; Redβ or RecT), enabling diverse, precise DNA manipulations ^1–6^. Owing to its operational simplicity and high efficiency, recombineering has been established as a powerful genome-editing platform ^7–11^. Beyond the model host *Escherichia coli*, recombineering have been deployed in a wide range of bacteria, including Gram-negative *Pseudomonas* ^12^, *Agrobacterium* ^13^, *Photorhabdus* ^14^, and *Burkholderia* ^15^, as well as Gram-positive *Bacillus* ^16^, *Lactobacillus* ^17^, *Mycobacterium* ^18^, and *Corynebacterium* ^19^, and have even been adapted for mammalian cells ^20^. Despite this broad host range, efficient recombineering tools remain underdeveloped for *Streptomyces* and other biotechnologically important actinobacteria, which are prolific producers of bioactive natural products.

Redα and RecE are both members of the PD-(D/E)XK phosphodiesterase superfamily, with distinct domain organizations: Redα contains a PD-EXK active site and a YqaJ domain, whereas RecE harbours a PD-DXK active site and a DUF3799 domain ^21, 22^. High-resolution structural studies have illuminated their distinct oligomeric architectures and catalytic mechanisms. The 2.4 Å crystal structure of Redα reveals a C3-symmetric toroidal homotrimer with a funnel-shaped central channel, which accommodates dsDNA at its wide opening and allows only ssDNA to pass through its narrow end ^23^. During processive dsDNA degradation, Redα associates with the linear undigested DNA strand in a beads-on-a-string fashion, progressing along the substrate until it reaches the DNA terminus or dissociates into monomers ^23^. A 1.88 Å crystal structure of the Redα–DNA complex further supports an electrostatic ratchet mechanism for processivity, in which forward translocation of Redα is driven by electrostatic attraction between the 5′-terminal phosphate of DNA and a positively charged pocket at the distal end of the active-site cleft ^24^. For *E. coli* RecE, a 2.8 Å crystal structure shows that it assembles into a C4-symmetric toroidal homotetramer with a tapered central channel: dsDNA enters via the wider opening, and ssDNA exits through the narrower end, which is partially occluded by the protein’s C-terminal segment ^25^. A 2.3 Å co-crystal structure of the Redα trimer in complex with three copies of the Redβ C-terminal domain (CTD) has defined the physical basis for recombinase coupling: each Redβ monomer binds the Redα trimer via its CTD, and Redα translocation along DNA sequentially loads Redβ monomers onto the nascent 3′ ssDNA overhang. The Redβ CTD also mediates interaction with the ssDNA-binding protein SSB, coordinating annealing of the 3′ overhang to SSB-coated dsDNA at the replication fork ^26^. While these studies have established the structural framework for prototypical RecE and Redα, the understanding of activity regulation in RecE/Redα exonucleases has remained largely confined to the core catalytic center, and the role of C-terminal non-catalytic sequence variation in modulating exonuclease activity and recombination performance remain poorly characterized.

CRISPR-Cas-based genome-editing technologies have greatly expanded the scope of bacterial genetic engineering ^27–36^. Nevertheless, achieving high-throughput, precise and efficient genome editing—especially for the integration of large DNA segments— remains a major challenge in many bacterial species. In *Streptomyces*, for example, CRISPR-Cas9-mediated genomic integration typically requires long homology arms and exhibits low efficiency for large DNA inserts, which limits its utility for rapid strain engineering. There is an urgent need to develop versatile, portable genome-editing tools to advance bacterial genome engineering capabilities.

In this study, we identify a panel of highly active actinobacteriophage RecET recombinase pairs that mediate efficient genome editing in *Streptomyces*, and systematically characterize the structural, biochemical and functional properties of Che9cRecE. We show that the extended C-terminal loop of Che9cRecE acts as a tunable regulatory module that calibrates exonuclease activity to optimize homologous recombination efficiency. We further demonstrate that this C-terminal module is portable across diverse phage exonucleases to boost recombineering performance, providing both mechanistic insights into the activity regulation of RecE/Redα family exonucleases and a generalizable strategy for advancing bacterial genome-editing technologies.

## Results

### Actinobacteriophage RecET pairs enable efficient homologous-recombination-mediated genome editing in *Streptomyces*

To identify RecET recombinases that enable efficient homologous-recombination-mediated genome editing in *Streptomyces*, we performed BLAST searches against the actinobacteriophage protein database to screen putative homologs of *E. coli* RecE and their cognate RecT pairing partners. After removing redundant sequences, we ultimately identified 31 RecET recombinase pairs (Supplementary Table 1).

To evaluate the genome-editing efficiency of these 31 actinobacteriophage RecET recombinase pairs in *Streptomyces*, we constructed plasmids designated pCas-actRecET-hyg (Fig. 1a) using pCRISPomyces2 as the backbone ^29^0. pCas-actRecET-hyg expresses CRISPR-Cas9, which introduces a double-stranded DNA break (DSB) at the *actVA5* locus of the *Streptomyces coelicolor* A3(2) chromosome; this DSB is a prerequisite for RecET recombinase-mediated homologous recombination. Each of the 31 RecET recombinase gene pairs was placed downstream of *cas9*, with both genes driven by the constitutive *rpsLp(XC)* promoter. Following homologous recombination mediated by actinobacteriophage RecET recombinases, the 1.9-kb hygromycin resistance cassette, flanked by 1,500-bp homology arms, integrates into the *actVA5* locus (Fig. 1a). We designated the genome editing method combining CRISPR-Cas9 with RecET recombinases as Cas-ET. We also built a control plasmid pCas-hyg (Fig. 1b) lacking RecET recombinase genes to compare editing efficiency against pCas-actRecET-hyg variants. Higher genome-editing efficiency for pCas-actRecET-hyg relative to pCas-hyg indicates that the corresponding RecET recombinases can mediate efficient homologous recombination in *S. coelicolor* A3(2).

**Fig. 1.**
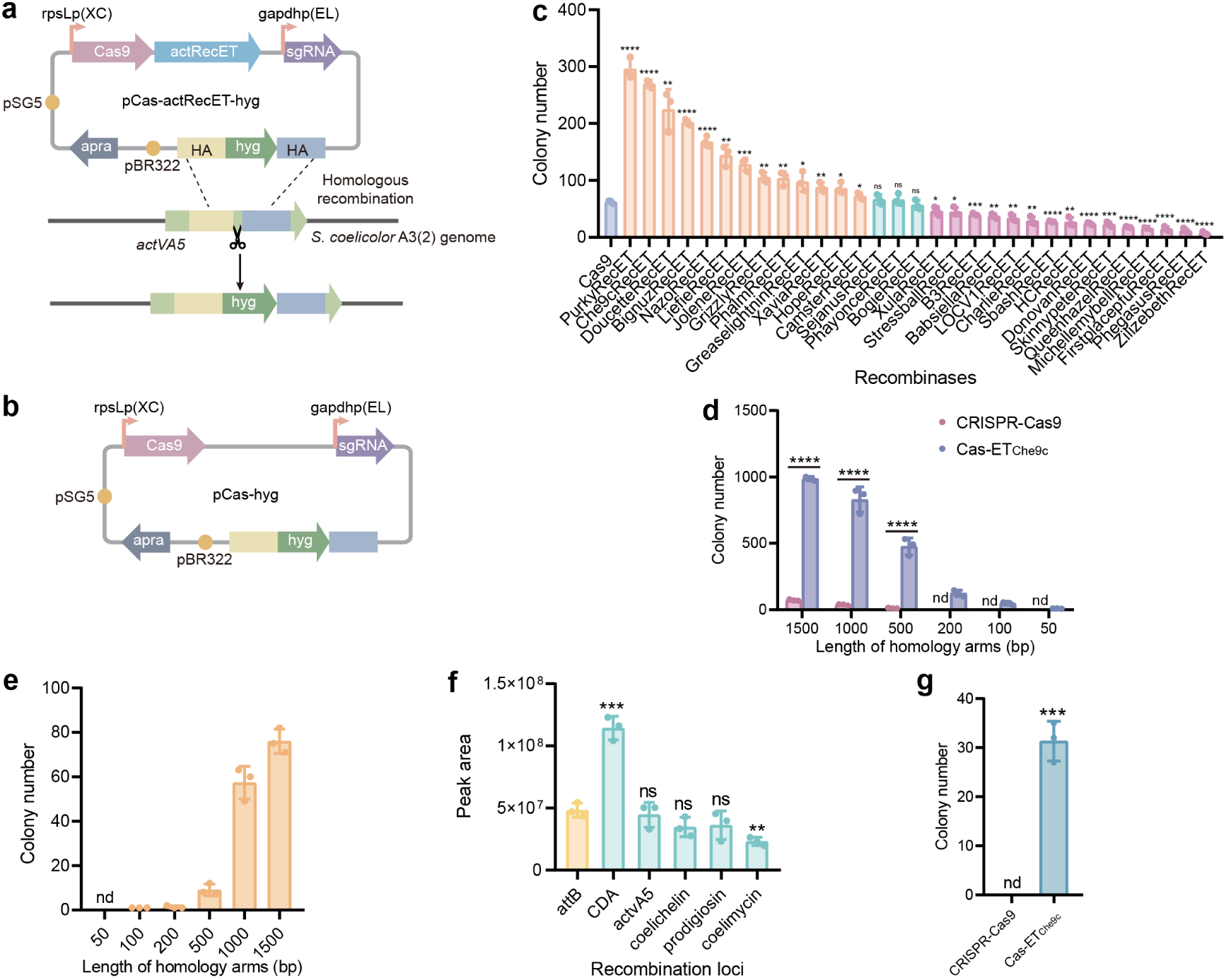
Actinobacteriophage-derived RecE homologs enable efficient homologous recombination-mediated genome editing in *Streptomyces*. **a,** Schematic of plasmid pCas-actRecET-hyg and insertion of the hygromycin resistance cassette at the *actVA5* locus of the *S. coelicolor* A3(2) chromosome. **b,** Schematic of plasmid pCas-hyg. **c,** Genome-editing efficiencies for insertion of the hygromycin resistance cassette flanked by 1,500-bp homology arms into the *actVA5* locus using CRISPR-Cas9 alone or Cas-ET. d, Genome-editing efficiencies for insertion of the hygromycin resistance cassette, flanked by homology arms of varying lengths, into the *actVA5* locus using CRISPR-Cas9 alone or Cas-ET_Che9c_. **e,** Genome-editing efficiencies for insertion of the 79-kb spinosyn biosynthetic gene cluster, flanked by homology arms of varying lengths, into the *actVA5* locus using Cas-ET_Che9c_. **f,** Spinosyn A yield from *S. coelicolor* A3(2) recombinant strains carrying the spinosyn biosynthetic gene cluster integrated at the indicated chromosomal loci. **g,** Genome-editing efficiencies for insertion of the hygromycin resistance cassette, flanked by 50-bp homology arms, into the *dptA* gene of the *S. roseosporus* NRRL 15998 chromosome using CRISPR-Cas9 alone or Cas-ET_Che9c_. Error bars represent mean ± s.d. of *n* = 3 biological replicates; dots denote values from individual replicates. Statistical significance was determined using two-tailed *t*-tests. \**P* ≤ 0.05, \*\**P* < 0.01, \*\*\**P* < 0.001, \*\*\*\**P* < 0.0001; ns, not significant; nd, not detected.

The results revealed that 13 RecET pairs are capable of mediating efficient homologous recombination in *S. coelicolor* A3(2). Among these, five RecET pairs originating from Che9c, Purky, Doucette, BigNuz and Nazo phages displayed the highest editing efficiencies (Fig. 1c). When PurkyRecET or Che9cRecET was combined with CRISPR-Cas9, the corresponding editing efficiencies were 4.8-fold and 4.4-fold higher, respectively, than that achieved with CRISPR-Cas9 alone (Fig. 1c).

We further compared the genome-editing efficiencies of Cas-ET_Che9c_ and CRISPR-Cas9 using homology arms of varying lengths: 50 bp, 100 bp, 200 bp, 500 bp, 1,000 bp, and 1,500 bp. The results showed that Cas-ET_Che9c_ required only 50-bp homology arms to insert the 1.9-kb hygromycin resistance cassette into the *actVA5* locus of *S. coelicolor* A3(2), whereas CRISPR-Cas9 required a minimum of 500-bp homology arms to accomplish the same integration (Fig. 1d). At homology arm lengths of 500 bp and 1,000 bp, the genome-editing efficiency of Cas-ET_Che9c_ was 43-fold and 22-fold higher, respectively, than that of CRISPR-Cas9 (Fig. 1d).

We also assessed the capacity of Cas-ET_Che9c_ to insert the 79-kb spinosyn biosynthetic gene cluster ^37^ into the *actVA5* locus of *S. coelicolor* A3(2), using homology arms of varying lengths: 50 bp, 100 bp, 200 bp, 500 bp, 1,000 bp, and 1,500 bp. The results revealed that Cas-ET_Che9c_ required only 100-bp homology arms to integrate the 79-kb spinosyn gene cluster into the *actVA5* locus of *S. coelicolor* A3(2) with 100% accuracy (Fig. 1e). In contrast, CRISPR-Cas9 failed to achieve this integration even with 1,500-bp homology arms.

We next tested Cas-ET_Che9c_ for insertion of the 79-kb spinosyn gene cluster into 5 additional loci distributed across the *S. coelicolor* A3(2) chromosome using 1,000-bp homology arms. These 5 loci are located in the calcium-dependent antibiotic (CDA) gene cluster, the coelichelin gene cluster, the prodigiosin gene cluster, the coelimycin gene cluster and the *phiC31-attB* site, respectively (Supplementary Fig. 1). The results demonstrated that Cas-ET_Che9c_ successfully inserted the 79-kb spinosyn gene cluster into all 5 tested loci in *S. coelicolor* A3(2) with 100% accuracy. Spinosyn A productions in these six *S. coelicolor* A3(2) recombinant strains harboring the spinosyn gene cluster were detected via high-performance liquid chromatography-mass spectrometry (HPLC-MS) analysis. The results revealed that all 6 recombinant strains successfully produced spinosyn A. Among them, the recombinant strain with the spinosyn gene cluster integrated at the CDA locus exhibited the highest spinosyn A yield, reaching 2.4-fold higher than that of the strain with integration at the *phiC31-attB* site — the most widely used locus for heterologous gene integration in *Streptomyces* (Fig. 1f).

We lastly tested Cas-ET_Che9c_ in a second *Streptomyces* strain, *S. roseosporus* NRRL 15998 — a producer of the antibacterial compound daptomycin (Supplementary Fig. 2). Using Cas-ET_Che9c_, we successfully inserted the 1.9-kb hygromycin resistance cassette into the *dptA* gene of the *S. roseosporus* NRRL 15998 genome with only 50-bp homology arms (Fig. 1g), whereas CRISPR-Cas9 alone could not achieve this integration with the same homology arms (Fig. 1g).

### Structural and sequence characterization of actinobacteriophage RecE homologs identifies a distinct Che9cRecE subfamily with unique C-terminal features

The protein 3-D structures of these 31 actinobacteriophage RecE homologs were predicted using AlphaFold3 ^38^. We next carried out multiple structural alignments (MSTA) of the 31 proteins together with *E. coli* RecE. From these alignments, we constructed similarity matrices that reflect the overall structural correlations among the proteins. These matrices were then used to build structural dendrograms via the unweighted pair-group method with arithmetic mean (UPGMA) ^39^. Protein clustering showed that 31 actinobacteriophage RecE homologs are hierarchically classified into several major structural clades, among which the high-efficiency actinobacteriophage RecE homologs (Che9CRecE, PurkyRecE, BigNuzRecE, NazoRecE) grouped within a distinct clade (Fig. 2a-b). Accordingly, we designated these four RecE homologs the Che9cRecE subfamily.

**Fig. 2.**
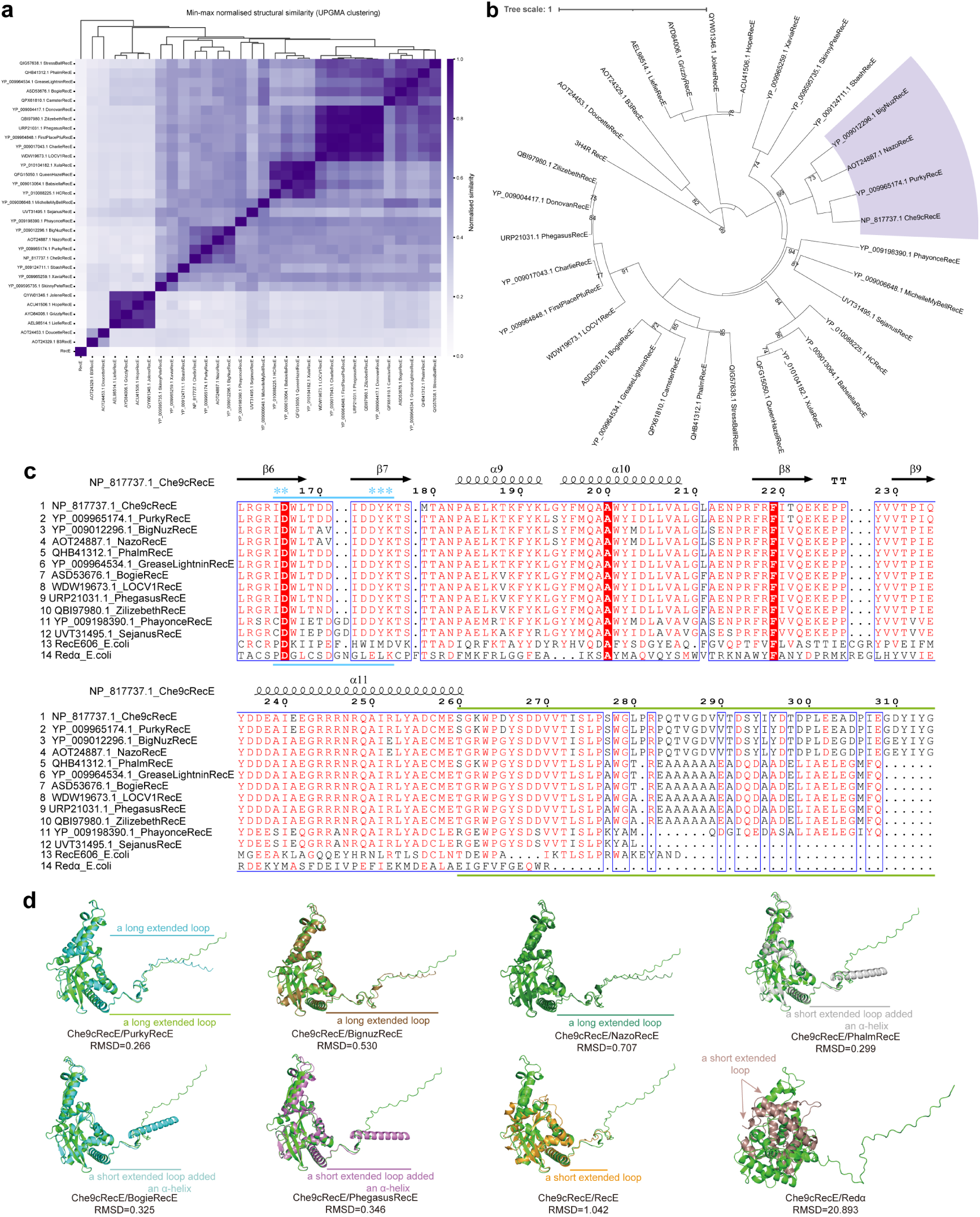
Structure and sequence features of actinobacteriophage RecE homologs. **a,** Structural dendrograms of RecE homologs. Heatmap color shading denotes the degree of similarity. **b,** Protein clustering of RecE homologs. Bootstrap values ≥70 are shown. **c,** Multiple sequence alignment of representative RecE homologs, showing conserved core regions of the PD-(D/E)XK superfamily and C-terminal regions. Secondary structural elements of Che9cRecE are annotated above the sequence alignment. The PD-(D/E)XK domains are marked with a blue line, and active-site residues are denoted by asterisks. The subfamily-specific C-terminal region of Che9cRecE is indicated by a green line. **d,** Structural comparison of AlphaFold3-predicted monomeric Che9cRecE and other RecE homologs, alongside experimentally determined *E. coli* RecE (PDB 3H4R) and Redα (PDB 1AVQ).

Protein domain analysis of the Che9cRecE subfamily (Che9CRecE, PurkyRecE, BigNuzRecE, NazoRecE) using the InterPro database revealed that these proteins contain PD-(D/E)XK-like (DUF3799) domains. Amino acid sequence alignment showed that members of the Che9cRecE subfamily harbour conserved ID-DYK active-site residues (Fig. 2c, Supplementary Fig. 3). Furthermore, their C-termini (the last 55 residues, residues 260–314 indicated by a green line) share sequence similarity and exhibit substantial divergence relative to other actinobacteriophage-derived RecE homologs, as well as *E. coli* RecE and Redα (Fig. 2c, Supplementary Fig. 3). The C-termini of Che9cRecE-subfamily proteins adopt unique long extended-loop structures (Fig. 2d, Supplementary Figs. 4 and 5). In contrast, the C-termini of other actinobacteriophage-derived RecE homologs form short extended loops appended with α-helical structures (Fig. 2d, Supplementary Figs. 4 and 5). The C-termini of *E. coli* RecE and Redα exhibit short extended-loop structures (Fig. 2d, Supplementary Fig. 4).

### Che9cRecE forms homotrimers and exhibits relatively low exonuclease activity

To determine the multimeric architecture of actinobacteriophage RecE homologs, we selected two representatives, Che9cRecE and BogieRecE, according to above sequence and structural characteristics. Che9cRecE is a 314-amino-acid protein with a monomer molecular weight of 35.6 kDa. BogieRecE is a 308-amino-acid protein with a monomer molecular weight of 34.5 kDa. N-terminally 6×His-tagged Che9cRecE and BogieRecE proteins were expressed in *E. coli* and purified to >95% purity (Fig. 3a). Size-exclusion chromatography indicated apparent molecular weights of 113.6 kDa for Che9cRecE (monomer mass: 35.6 kDa) and 104.4 kDa for BogieRecE (monomer mass: 34.5 kDa) (Fig. 3b), demonstrating that both proteins form homotrimers in solution. This result indicates that the homotrimeric architecture is conserved among actinobacteriophage RecE homologs.

**Fig. 3.**
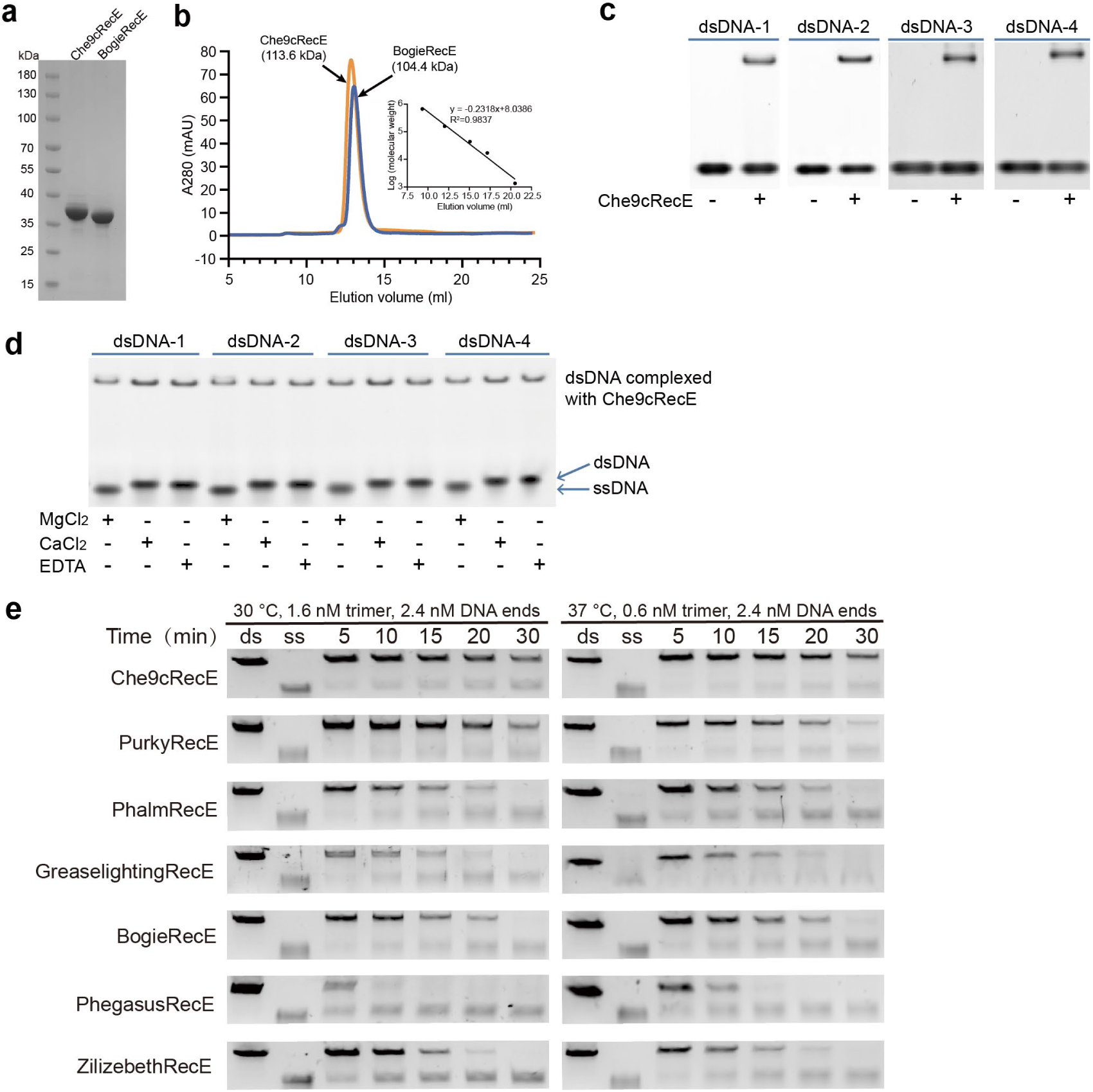
The multimeric architecture and exonuclease activity of actinobacteriophage RecE homologs. **a,** SDS–PAGE of purified proteins Che9cRecE and BogieRecE. **b,** Size-exclusion chromatography profiles of Che9cRecE (orange) and BogieRecE (blue). **c,** EMSA of Che9cRecE with four dsDNA substrates. Che9cRecE and dsDNA were mixed at a 1:1 molar ratio. **d,** Agarose gel electrophoresis analysis of products from Che9cRecE exonuclease activity assays performed in reaction buffers containing MgCl_2_, CaCl_2_, or EDTA. **e,** Agarose gel analysis of products from exonuclease activity assays of RecE homologs. Aliquots harvested at 5, 10, 15, 20, or 30 min post-reaction were analyzed. Lanes ds and ss correspond to the dsDNA substrate and ssDNA product, respectively.

To examine the binding of Che9cRecE to dsDNA, we synthesized two sets of oligonucleotides whose sequences were derived from the homology arms used for inserting the hygromycin resistance cassette at the *actVA5* locus of *S. coelicolor* A3(2). Each set comprised two 17-mer ssDNA oligos bearing either a 5′-OH or 5′-PO₄ terminus, alongside their complementary 15-mer oligo with a 5′-OH end. Each set of strands were pairwise annealed to generate 15 bp DNA duplexes containing a 5′-hydroxylated (dsDNA-1, dsDNA-3) or 5′-phosphorylated (dsDNA-2, dsDNA-4) dinucleotide overhangs.

Protein electrophoretic mobility shift assays (EMSA) showed that Che9cRecE–dsDNA complex bands were detectable at a 1:1 molar ratio of Che9cRecE protein to dsDNA, confirming direct binding between Che9cRecE and dsDNA. No significant difference was observed in the band patterns formed between Che9cRecE and the four dsDNA substrates bearing either 5′-OH or 5′-PO₄ termini (Fig. 3c). Che9cRecE was individually incubated with dsDNA-1 to dsDNA-4 at a 1:1 molar ratio in reaction buffers supplemented with MgCl_2_, CaCl_2_, or EDTA. Gel electrophoresis analysis revealed that Che9cRecE possesses exonuclease activity in MgCl_2_-containing buffer, where it processively digests dsDNA to generate ssDNA. In contrast, Che9cRecE exhibited no exonuclease activity in CaCl_2_ buffer, consistent with the abolished activity observed when Mg²⁺ was chelated by EDTA. Likewise, no obvious activity difference was detected for Che9cRecE acting on the four dsDNAs with 5′-OH versus 5′-PO₄ termini (Fig. 3d).

To investigate the mechanism underlying the higher homologous recombination efficiency of Che9cRecET and PurkyRecET, we characterized the exonuclease activity of Che9cRecE and PurkyRecE and compared it with that of several representative actinobacteriophage RecE homologs. In this assay, we monitored the digestion of linear 2,686-bp pUC19 dsDNA using enzyme concentrations lower than that of the DNA ends. Agarose gel electrophoresis with SYBR Gold staining was used to quantify the remaining dsDNA substrate and the ssDNA products generated during the reaction. We selected two assay temperatures: 37 °C, the growth temperature of *Mycobacterium* (the original host used for phage isolation), and 30 °C, the cultivation temperature of *Streptomyces*, the genus used for genome editing by these RecE homologs. We therefore measured the exonuclease activity of representative RecE homologs at both temperatures. The results revealed that Che9cRecE and PurkyRecE displayed weaker exonuclease activity relative to RecE homologs with moderate or low editing efficiency (PhalmRecE, GreaseLightninRecE, BogieRecE, PhegasusRecE, ZilizebethRecE), when tested using 2.4 nM DNA ends and 1.6 nM trimeric enzymes at 30 °C and 0.6 nM trimeric enzymes at 37 °C (Fig. 3e).

These results reveal that Che9cRecE and PurkyRecE exhibit relatively low exonuclease activity. Therefore, we speculated that the high gene editing efficiencies of Che9cRecET and PurkyRecET might be attributed to low exonuclease activity of Che9cRecE and PurkyRecE.

### Cryo-EM structure of Che9cRecE reveals its toroidal homotrimer

We further determined the cryo-EM structure of Che9cRecE. Briefly, Che9cRecE was heterologously expressed in *Escherichia coli* and purified sequentially using HisTrap HP, HiTrap Q FF, and Superdex 200 Increase 10/300 GL columns. The purified protein was plunge-frozen for subsequent cryo-EM data collection. A total of 4,979 micrographs were acquired, and 3,521,633 particles were extracted and curated using cryoSPARC ^40^. Micrograph analysis revealed that the Che9cRecE monomer self-assembles into a C3-symmetric toroidal homotrimer, with a distinct central cavity observed in the trimeric architecture (Fig. 4a and 4b). This overall structural fold is highly analogous to that of the previously reported *E. coli* Redα homotrimer ^23^.

**Fig. 4.**
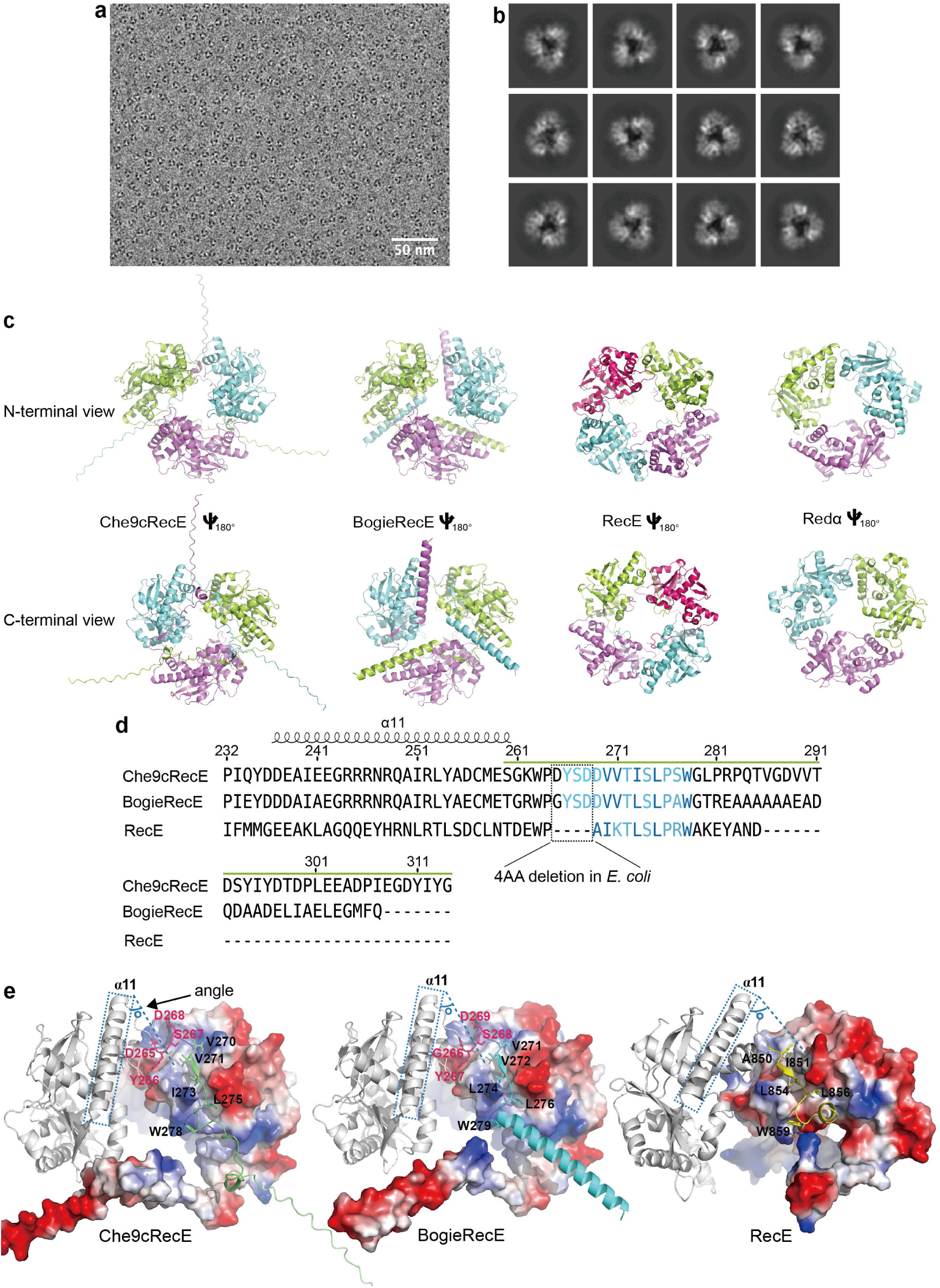
Protein structures of Che9cRecE and BogieRecE, and their comparison with *E. coli* RecE and Redα. **a,** Representative 2D class from cryo-EM particle classification of Che9cRecE. **b,** Example cryo-EM micrograph of Che9cRecE collected at 105,000× magnification, selected from 4,979 micrographs with consistent features. **c,** AlphaFold3-predicted homotrimer models of Che9cRecE and BogieRecE, alongside the experimentally determined homotetramer structure of *E. coli* RecE (PDB 3H4R) and homotrimer of Redα (PDB 1AVQ). **d**, Sequence alignment of the C-terminal regions of Che9cRecE, BogieRecE and *E. coli* RecE. The C-terminal loop region of Che9cRecE is marked with a green line. Residues located at inter-subunit interfaces are colored blue; hydrophobic core residues are highlighted in bold. **e**, Inter-subunit interface diagrams for Che9cRecE, BogieRecE and *E. coli* RecE. The angles between α-helix 11 and the β-sheet immediately following it were shown. The blue dashed lines are parallel to this β-sheet. For *E. coli* RecE, the AlphaFold3-predicted tetramer structure was used because its experimental structure does not include the last two C-terminal residues. The DYSD motif of Che9cRecE and GYSD motif of BogieRecE are in magenta. Hydrophobic core residues interacting with the adjacent subunit are shown in black.

In total, 4,207,765 qualified particles were subjected to multiple rounds of 2D classification, followed by iterative homogeneous and non-uniform refinements, yielding a final 3D reconstruction at an estimated resolution of 4.35 Å (Supplementary Fig. 6). The central cavity of the Che9cRecE homotrimer forms a funnel-shaped channel, which shares high structural similarity with the channel of *E. coli* Redα (Supplementary Fig. 6) ^23^. Notably, ambiguous density blobs were detected beneath the Che9cRecE trimer, which were speculated to correspond to the C-terminal regions of Che9cRecE (Supplementary Fig. 6).

Structural docking of the AlphaFold3-predicted Che9cRecE model into our experimental 3D density map showed that only the N-terminal region (residues 1–259) could be reliably fitted into the map. The unassigned peripheral density was therefore hypothesized to arise from the Che9cRecE C-terminus (residues 260–314), which is predicted by AlphaFold3 to form an extended flexible loop (Supplementary Fig. 6). Of note, preferred particle orientation limited the achievement of a higher-resolution map despite C3 symmetry imposition. Although the current resolution restricts atomic-level structural interpretation, our cryo-EM reconstructions unambiguously confirm that Che9cRecE adopts a homotrimeric assembly (Fig. 4a, 4b; Supplementary Fig. 6). This trimeric organization represents a structural divergence from the canonical homotetrameric architecture of *E. coli* RecE ^25^.

Given the limited resolution of our cryo-EM reconstruction, which prevented detailed atomic-level structural interpretation, we used AlphaFold3 to generate homotrimeric models of Che9cRecE and BogieRecE (Fig. 4c). The predicted assemblies of Che9cRecE (ipTM = 0.83, pTM = 0.85) and BogieRecE (ipTM = 0.83, pTM = 0.84) returned high-confidence scores. AlphaFold3 predictions showed that both Che9cRecE and BogieRecE assemble into C3-symmetric toroidal homotrimers. Che9cRecE and BogieRecE trimers share highly similar overall folds; their most prominent structural difference lies at the C-terminus: Che9cRecE bears a C-terminal long extended-loop, whereas BogieRecE contains a C-terminal short extended loop appended with an α-helix (Fig. 4c; also Fig. 2d). Previous work established that *E. coli* RecE forms a C4-symmetric toroidal homotetramer ^25^, while Redα adopts a C3-symmetric toroidal homotrimer ^23^. Comparison of our AlphaFold3 models for Che9cRecE and BogieRecE with the experimentally solved structures of *E. coli* RecE homotetramer (PDB 3H4R) and Redα homotrimer (PDB 1AVQ) showed that the oligomeric states of Che9cRecE and BogieRecE are more closely related to Redα than to *E. coli* RecE. Notably, the Che9cRecE monomer fold resembles that of RecE more strongly than Redα (Fig. 2d). Collectively, the Che9cRecE and BogieRecE trimers define a distinct class of exonuclease trimers within the Red/ET system, alongside Redα.

We next compared the homomultimeric architectures of Che9cRecE, BogieRecE, and *E. coli* RecE. At the C-terminal regions, residues 266–278 of Che9cRecE, residues 267–279 of BogieRecE, and residues 850–859 of *E. coli* RecE (blue in Fig. 4d) reside at inter-subunit interfaces (Fig. 4e). Hydrophobic residues V270, V271, I273, L275 and W278 of Che9cRecE; V271, V272, L274, L276 and W279 of BogieRecE; and A850, I851, L854, L856, and W859 of *E. coli* RecE (bold blue in Fig. 4d, black in Fig. 4e) form the hydrophobic core that mediates subunit contacts. Sequence alignment revealed that the DYSD motif of Che9cRecE and GYSD motif of BogieRecE (boxed in Fig. 4d; magenta in Fig. 4e) located downstream of α-helix 11 at the inter-subunit interfaces are absent in *E. coli* RecE. Quantification of the angles between α-helix 11 and the β-sheet immediately following it showed that these angles are smaller in the Che9cRecE and BogieRecE homotrimers relative to those in the *E. coli* RecE homotetramer (Fig. 4e). These observations demonstrate distinct inter-subunit packing geometry between the Che9cRecE/BogieRecE trimers and the *E. coli* RecE tetramer. We hypothesize that the lack of these four residues in *E. coli* RecE remodels the inter-subunit packing angle and acts as a key determinant governing the different oligomeric states. The inter-subunit interface of Redα shares very low similarity with those of Che9cRecE, BogieRecE, and *E. coli* RecE (Supplementary Fig. 7), suggesting that the Redα homotrimer adopts an assembly mechanism distinct from that of the Che9cRecE/BogieRecE homotrimers.

### DNA binding and electrostatic functional polarization of Che9cRecE and BogieRecE

To investigate how Che9cRecE and BogieRecE engage DNA, we used AlphaFold3 to model their complexes with a partial-duplex DNA substrate containing a double-stranded region and single-stranded overhang. The two models adopted highly similar architectures and resembled the homotetramer of *E. coli* RecE and homotrimer of Redα reported previously ^23–25^. In these models, DNA resides inside the funnel-shaped central channel of the homotrimers. The N-terminal regions of the assembly form a wide entrance channel for dsDNA, while the C-termini converge to form a narrow central pore through which ssDNA exits (Fig. 5a).

**Fig. 5.**
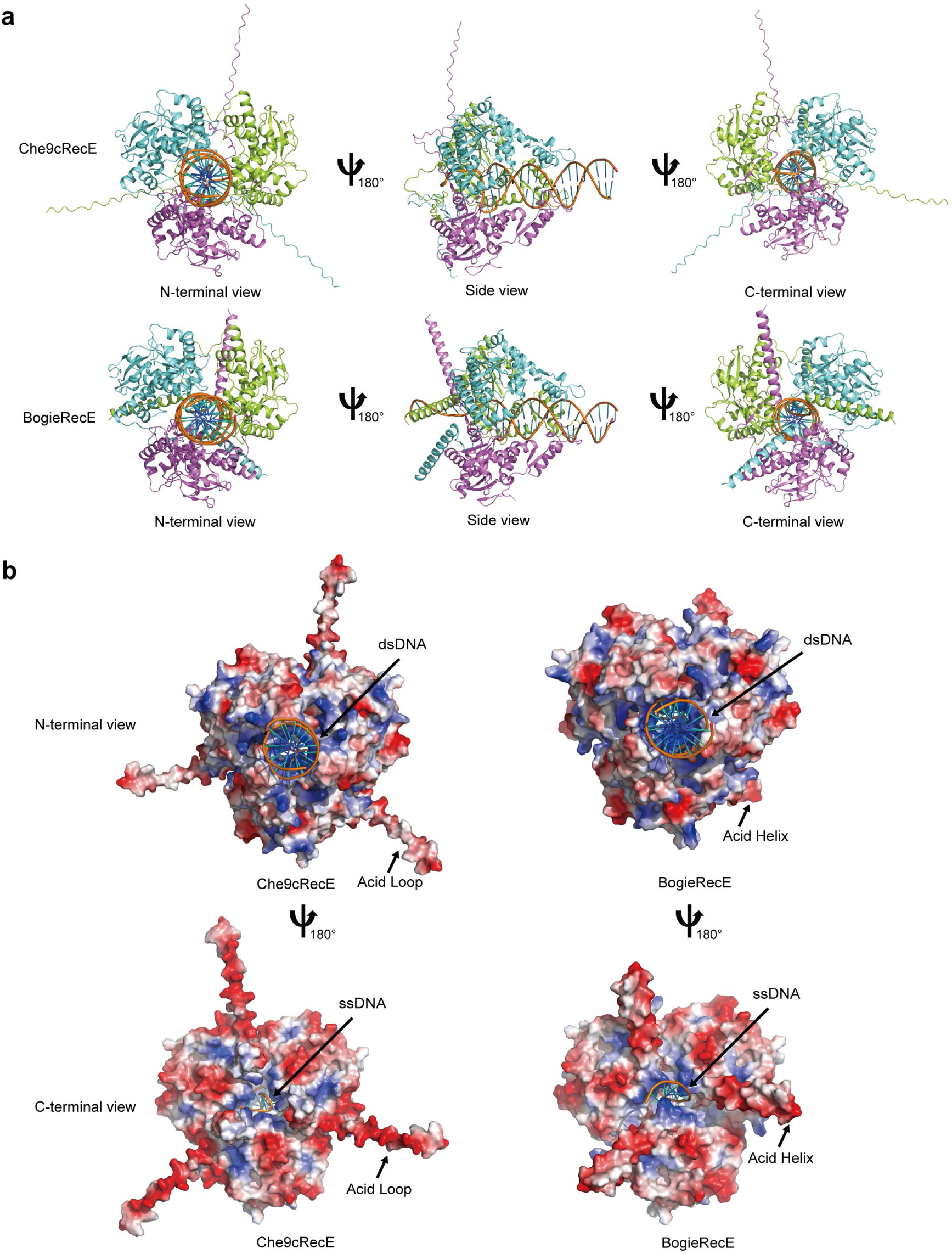
AlphaFold3-modelled homotrimer structures of Che9cRecE and BogieRecE in complex with DNA. **a,** AlphaFold3-predicted homotrimer structures of Che9cRecE and BogieRecE in complex with a partial duplex DNA substrate. Structures are displayed from the N-terminal, side and C-terminal perspectives, respectively. **b,** Electrostatic surface representations of Che9cRecE and BogieRecE in complex with a partial duplex DNA substrate; negative electrostatic potential is coloured red and positive potential blue. Views are shown from the N-terminus and C-terminus, respectively.

The Che9cRecE–DNA and BogieRecE–DNA complex models also show nearly identical DNA-binding modes at the residue level. In both complexes, positively charged arginine and lysine residues, R282, K187, K192, K53, K54, K92, K84 and R100 in Che9cRecE and R282, K188, K193, K54, K55, K93, K85 and R101 in BogieRecE, contributed by all three subunits converged on the DNA duplex and formed an extensive salt-bridge network with the DNA phosphate backbone (Supplementary Fig. 8). These basic residues lining the central channel at equivalent positions in both complexes supports a conserved DNA-binding mode.

Electrostatic surface analysis revealed that both Che9cRecE and BogieRecE homotrimers display marked functional polarization of their central channel. The wide entry portals of these funnel-shaped homotrimers possess strong positive electrostatic potential, generating an electropositive surface that captures and directs negatively charged dsDNA substrates into the central channel (Fig. 5b). By contrast, the base of the funnel assembly, particularly surrounding the narrow central pore, features a negatively charged ring formed mainly by acidic residues within the C-terminal region (Fig. 5b). Collectively, these observations support a bidirectional electrostatic mechanism: the positively charged wide entry portals facilitate capture and loading of dsDNA substrates, whereas the negatively charged base repels the newly generated ssDNA product, establishing an electrostatic gradient that drives processive 5′–3′ exonucleolytic activity.

Notably, the Che9cRecE homotrimer contains a flexible C-terminal loop, whereas the BogieRecE homotrimer harbours a structured C-terminal helix; both elements lie adjacent to the ssDNA exit pore (Fig. 5b). This structural difference raises the possibility that divergent C-terminal conformations among RecE homologs may differentially modulate DNA release dynamics during DNA processing.

### Effects of C-terminal substitution of actinobacteriophage RecE homologs on exonuclease activity and genome editing efficiency

The 55-residue Che9cRecE C-terminal loop (Che9cRecECL) region (residues 260–314, green in Fig. 2c) contains residues 266–278 that mediate subunit contacts; the remaining 42 residues may primarily be involved in modulating DNA release dynamics during DNA processing (Fig. 4d). To dissect how Che9cRecECL regulates exonuclease activity and genome-editing efficiency, we individually replaced the native short C-terminal extended loops with appended α-helical structures (Fig. 2d) from four actinobacteriophage RecE homologs (PhalmRecE, BogieRecE, PhegasusRecE, ZilizebethRecE). These four homologs exhibited high exonuclease activity yet low genome-editing efficiency (Fig. 3e). We constructed plasmids pCas-actRecEΔC-Che9cRecECL-RecT (Fig. 6a) to express chimeric RecE variants together with their cognate RecT partners. These chimeric pairs were incorporated into the Cas-ET genome-editing system for insertion of a 1.9-kb hygromycin resistance cassette at the *actVA5* locus of the *S. coelicolor* A3(2) chromosome.

**Fig. 6.**
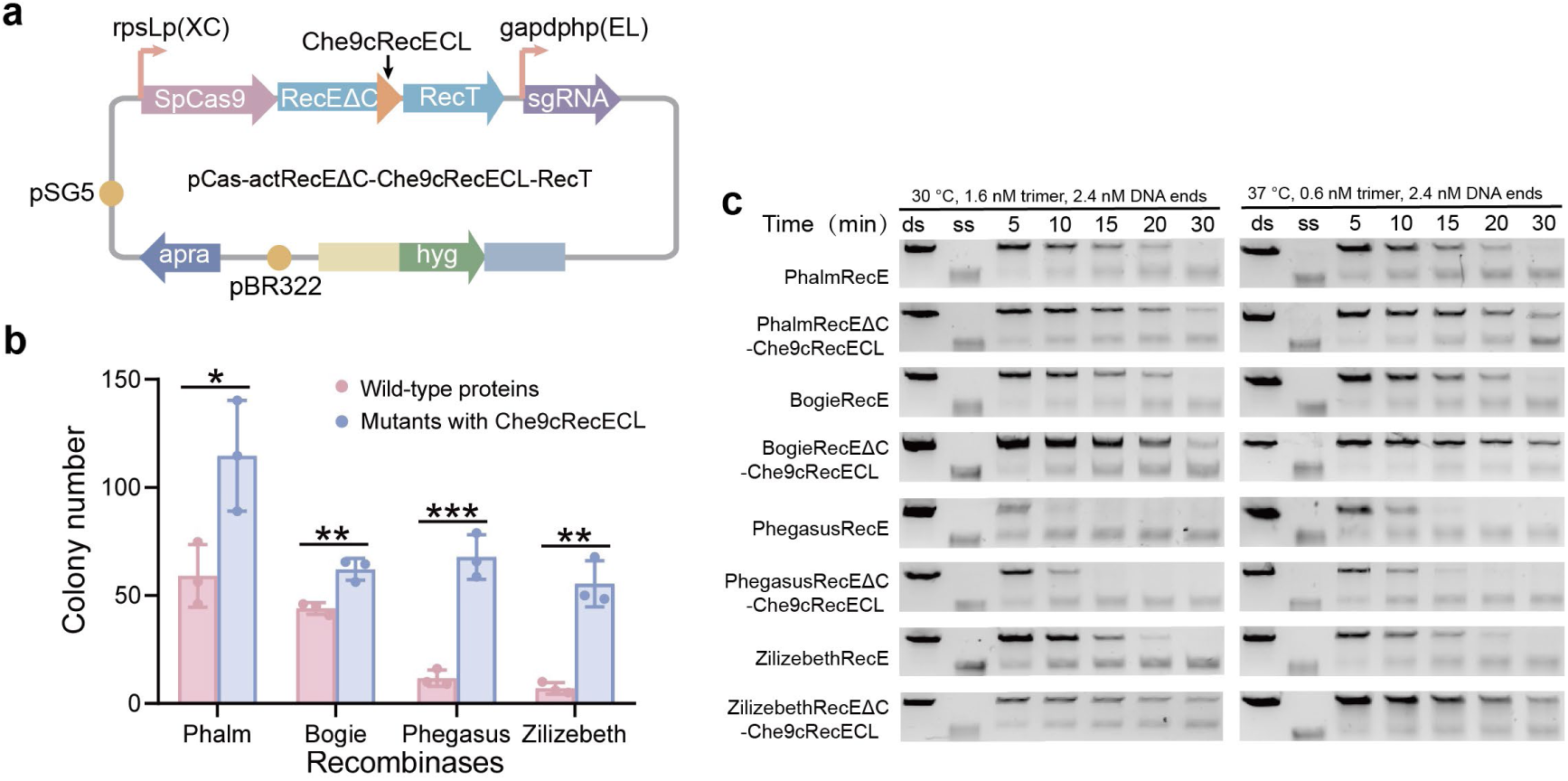
Effects of C-terminal substitution of actinobacteriophage RecE homologs on exonuclease activity and genome editing efficiency. **a,** Schematic diagram of plasmid pCas-actRecEΔC-Che9cRecECL-RecT. **b,** Genome-editing efficiencies for insertion of a hygromycin resistance cassette flanked by 1,500-bp homology arms into the *actVA5* locus of *S. coelicolor* via the Cas-ET system. Each actinobacteriophage RecT was combined with its cognate wild-type RecE or chimeric RecE variant, in which the native C-terminus was individually replaced with Che9cRecECL. Error bars represent mean ± s.d. of *n* = 3 biological replicates; dots indicate values from individual replicates. Statistical significance was determined using two-tailed *t*-tests. \**P* < 0.05, \*\**P* < 0.01, \*\*\**P* < 0.001. **c,** Exonuclease activity assays of wild-type actinobacteriophage RecE proteins and chimeric RecE variants in which native C-termini were individually replaced with Che9cRecECL. Aliquots collected at indicated time post-reaction were analyzed. Lanes ds and ss correspond to the dsDNA substrate and ssDNA product, respectively.

Genome-editing assays showed that all four chimeric RecE proteins conferred substantially higher editing efficiency in *S. coelicolor* A3(2) compared with their wild-type counterparts. Notably, the Phegasus and Zilizebeth chimeras yielded 5.7-fold and 7.9-fold improvements, respectively (Fig. 6b).

Exonuclease activity assays revealed that replacement of the native C-termini of PhalmRecE, BogieRecE, PhegasusRecE and ZilizebethRecE with Che9cRecECL markedly reduced the exonuclease activity of the resulting chimeras compared with their wild-type counterparts (Fig. 6c). Together, these results demonstrate that Che9cRecECL modulates genome-editing efficiency by tuning exonuclease activity.

PhalmRecE, BogieRecE, PhegasusRecE and ZilizebethRecE share high sequence conservation with Che9cRecE at the C-terminal inter-subunit interface (Figs. 2c and Supplementary Fig. 9). We hypothesize that these proteins adopt a similar multimeric assembly mechanism, and that swapping their native C-termini for Che9cRecECL does not impair multimerization. Thus, Che9cRecECL is portable across divergent actinobacteriophage RecE homologs for modulation of their exonuclease activity and genome-editing performance.

### Fusion of Che9cRecECL to the C-terminal ends of *E. coli* Redα, *E. coli* RecE, and *Pseudomonas* phage exonuclease A enhances their homologous-recombination efficiencies

We assessed the portability of Che9cRecECL for modulating exonuclease activity and homologous-recombination efficiency across a wider panel of bacteriophage exonucleases beyond actinobacteriophage-derived RecE homologs. We first replaced the native 11-residue C-terminus of *E. coli* Redα (green in Fig. 2c) with Che9cRecECL. The gene encoding this chimeric RedαΔC-Che9cRecECL protein was paired with its cognate Redβ gene to generate plasmid pSC101-BAD-gbaΔC-Che9cRecECL-tet (Supplementary Fig. 10a). Linear-circular recombineering efficiency mediated by pSC101-BAD-gbaΔC-Che9cRecECL-tet was compared with that of pSC101-BAD-gba-tet carrying the wild-type Redα coding sequence in *E. coli* GB2005 (Supplementary Fig. 10b). The results revealed that the chimeric RedαΔC-Che9cRecECL protein exhibited 33.5-fold lower linear-circular recombineering efficiency relative to wild-type Redα (Supplementary Fig. 10c). We hypothesized that this substantial reduction in recombineering efficiency for RedαΔC-Che9cRecECL arose from fundamentally distinct assembly mechanisms of the Redα homotrimer and Che9cRecE homotrimer. Substitution of the native 11-residue C-terminus of Redα with Che9cRecECL may disrupt Redα homotrimer assembly, consequently diminishing recombineering efficiency.

In an alternative strategy, Che9cRecECL was directly fused to the C-terminal end of Redα to generate the chimeric Redα-Che9cRecECL protein. The gene encoding this chimeric Redα variant was paired with its cognate Redβ gene to construct plasmid pSC101-BAD-gba-Che9cRecECL-tet (Fig. 7a). Linear-circular recombineering (Fig. 7b) efficiency mediated by pSC101-BAD-gba-Che9cRecECL-tet was compared with that of pSC101-BAD-gba-tet in *E. coli* GB2005. The results revealed that the chimeric Redα-Che9cRecECL protein exhibited a 2.8-fold higher linear-circular recombineering efficiency relative to wild-type Redα (Fig. 7c). Exonuclease activity of Redα-Che9cRecECL was significantly lower than that of wild-type Redα at 37 °C using 0.3 nM or 0.6 nM trimeric enzyme (Supplementary Fig. 11). These findings further demonstrated that Che9cRecECL can modulate gene-editing efficiency by regulating exonuclease activity.

**Fig. 7.**
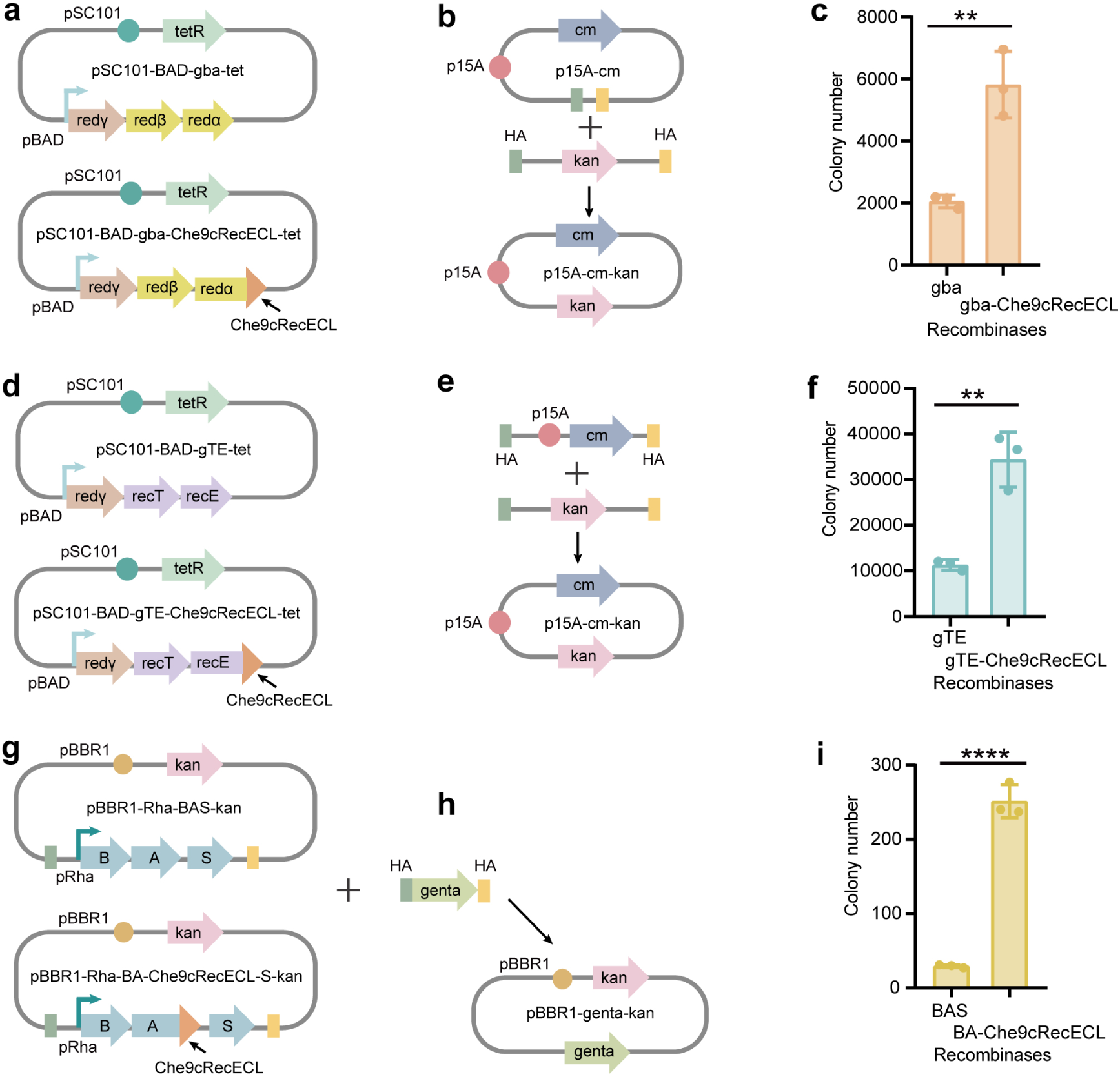
Fusion of Che9cRecECL to the C-terminal ends of bacteriophage exonucleases enhances their recombineering efficiencies across diverse bacterial species. **a,** Schematic diagrams of plasmids pSC101-BAD-gba-tet and pSC101-BAD-gba-Che9cRecECL-tet. **b,** Schematic diagram of linear-circular recombineering. **c,** Linear-circular recombineering efficiency mediated by pSC101-BAD-gba-tet and pSC101-BAD-gba-Che9cRecECL-tet in *E. coli* GB2005. **d,** Schematic diagrams of plasmids pSC101-BAD-gTE-tet and pSC101-BAD-gTE-Che9cRecECL-tet. **e,** Schematic diagram of linear-linear recombineering. **f,** Linear-linear recombineering efficiency mediated by pSC101-BAD-gTE-tet and pSC101-BAD-gTE-Che9cRecECL-tet in *E. coli* GB2005. **g,** Schematic diagrams of plasmids pBBR1-Rha-BAS-kan and pBBR1-Rha-BA-Che9cRecECL-S-kan. **h,** Schematic diagram of linear-circular recombineering. **i,** Linear-circular recombineering efficiency mediated by pBBR1-Rha-BAS-kan and pBBR1-Rha-BA-Che9cRecECL-S-kan in *P. putida* KT2440. Error bars represent mean ± s.d. of *n* = 3 biological replicates; dots indicate values from individual replicates. Statistical significance was determined using two-tailed *t*-tests. \*\**P* < 0.01, \*\*\*\**P* < 0.0001.

Che9cRecECL was also directly fused to the C-terminal end of *E. coli* RecE (EcRecE) to generate the chimeric EcRecE-Che9cRecECL protein. The gene encoding this chimeric EcRecE-Che9cRecECL protein was paired with its cognate EcRecT gene to construct plasmid pSC101-BAD-gTE-Che9cRecECL-tet (Fig. 7d). Linear-linear recombineering (Fig. 7e) efficiency mediated by pSC101-BAD-gTE-Che9cRecECL-tet was compared with that of pSC101-BAD-gTE-tet, which carries the wild-type EcRecE coding sequence in *E. coli* GB2005. The results revealed that the chimeric EcRecE-Che9cRecECL protein exhibited a 3-fold higher linear-linear recombineering efficiency relative to wild-type EcRecE (Fig. 7f).

Che9cRecECL was further directly fused to the C-terminal end of the exonuclease A from *Pseudomonas* phage Ab31 ^12^ to generate the chimeric A-Che9cRecECL protein. The gene encoding this chimeric A-Che9cRecECL protein was paired with its cognate SSAP B gene to construct plasmid pBBR1-Rha-BA-Che9cRecECL-S-kan (Fig. 7g). Linear-circular recombineering (Fig. 7h) efficiency mediated by pBBR1-Rha-BA-Che9cRecECL-S-kan was compared with that of pBBR1-Rha-BAS-kan carrying the wild-type A coding sequence in *Pseudomonas putida* KT2440. The results revealed that the chimeric A-Che9cRecECL protein exhibited an 8.7-fold higher linear-circular recombineering efficiency relative to wild-type A (Fig. 7i).

## Discussion

This study addresses a major limitation of current *Streptomyces* genetic tools, which typically require long homology regions and suffer from low integration efficiency for large biosynthetic gene clusters. We establish Cas-ET, a CRISPR-Cas9-assisted, actinobacteriophage-derived RecET system that enables highly efficient homologous recombination-mediated genome editing in *Streptomyces*. We identified 13 naturally occurring RecET pairs with superior editing performance, among which Che9cRecET stands out by enabling integration of a 1.9-kb fragment with homology arms as short as 50 bp. By comparison, CRISPR-Cas9 alone required a minimum of 500-bp homology arms to achieve the same integration. With 500-bp homology arms, Cas-ET_Che9c_ yielded 42-fold higher genome-editing efficiencies relative to CRISPR-Cas9 (Fig. 1d). Furthermore, Cas-ET_Che9c_ enabled accurate integration of large DNA fragments >75 kb using merely 100-bp homology arms across multiple chromosomal loci (Fig. 1e and f). In contrast, CRISPR-Cas9 could not accomplish this integration even with 1 500-bp homology arms.

Our study reveals an unanticipated relationship: higher intrinsic exonuclease activity among RecE homologs consistently correlates with diminished genome-editing performance (Fig. 3e and Fig. 6c). Consistent with this trend, the two most efficient recombinases, Che9cRecE and PurkyRecE, display substantially weaker exonuclease activity than homologs with moderate or low editing efficiency (Fig. 3e). These observations support a model in which homologous recombination efficiency is governed by an optimal window of exonuclease activity. Excessive exonuclease activity produces overly extended 3′ ssDNA overhangs prone to intramolecular secondary structure formation, which in turn reduces the effective length of homologous sequence competent to support stable RecT-DNA filament assembly and productive strand pairing. Conversely, insufficient exonuclease activity also fails to generate 3′ ssDNA overhangs of sufficient length to support RecT-DNA filament assembly and productive strand pairing. Che9cRecE appears evolutionarily tuned to reside within this optimal activity window, generating 3′ overhangs of sufficient length to drive efficient recombination while avoiding the detrimental effects of excessive 5′ resection.

The Che9cRecECL is essential for maintaining exonuclease activity within the optimal range. Notably, swapping the native C-termini of four low-efficiency RecE homologs with Che9cRecECL consistently reduced their exonuclease activities and conferred substantial improvements in genome-editing efficiency, with enhancements of up to 7.9-fold relative to their wild-type counterparts (Fig. 6b). These gain-of-function results define Che9cRecECL as a portable regulatory module that can re-calibrate the catalytic output of diverse RecE exonucleases. Given that these low-efficiency homologs share high sequence conservation at the inter-subunit interface with Che9cRecE, we infer that C-terminal substitution does not perturb trimeric assembly.

Within the conserved mechanistic framework of bidirectional electrostatic model of RecE family homomultimers, subtle differences in C-terminal conformation — an extended flexible loop in high-efficiency Che9cRecE versus a rigid α-helix in low-efficiency BogieRecE — fine-tune exonuclease activity. We hypothesize that the flexible C-terminal loop, positioned adjacent to the ssDNA exit pore, introduces dynamic steric occlusion that slows ssDNA egress and limits processivity, resulting in lower overall exonuclease activity. In contrast, the rigid C-terminal helix of less active variants permits unimpeded DNA release and sustained digestion, leading to higher exonuclease activity output but diminished recombination efficiency. This structure-function relationship provides a molecular explanation for how C-terminal sequence divergence translates into dramatic differences in genome-editing performance. Prior to this work, the understanding of how RecE exonucleases are regulated at the activity level had largely been restricted to the core catalytic layer, encompassing fundamental functional features such as catalytic residues in the active site, electrostatic interactions mediating DNA binding, and the electrostatic ratchet mechanism that drives enzyme processivity ^23–26^. The C-terminal loop identified in this study forms a fine-tuning regulatory structure separate from the core catalytic center. Instead of directly participating in the hydrolysis of DNA phosphodiester bonds, it calibrates exonuclease activity to an optimal window for homologous recombination by controlling the release rate of ssDNA and limiting the enzyme’s processive digestion capacity.

The broader utility of Che9cRecECL as an engineering module is validated by direct fusion of Che9cRecECL to the C-termini of three phylogenetically distinct phage exonucleases — *E. coli* Redα, *E. coli* RecE, and *Pseudomonas* phage Ab31 exonuclease A — uniformly enhanced recombineering efficiency by 2.8- to 8.7-fold in their respective bacterial hosts (Fig. 7). Notably, this improvement was observed for both trimeric (Redα) and tetrameric (RecE) exonucleases. Previous studies have demonstrated that Redαβ mediates efficient genome editing across more than ten bacterial species, spanning Gamma-proteobacteria (e.g., *E. coli*, *Shigella flexneri*, *Salmonella enterica*, *Pseudomonas aeruginosa*, *Vibrio cholerae* and *Klebsiella pneumoniae*) ^2, 41–49^, Beta-proteobacteria (e.g., *Burkholderia cepacia*, *Burkholderia thailandensis* and *Burkholderia pseudomallei*) ^15, 50^, and Alpha-proteobacteria (e.g., *Agrobacterium tumefaciens*) ^51^. Combined with our experimental evidence, these findings establish C-terminal fusion as a general, robust strategy for engineering diverse exonucleases to boost recombineering efficiency.

Our structural and biochemical characterization further reveals that actinobacteriophage RecE proteins form a distinct clade of C3-symmetric toroidal homotrimers, representing a new architectural class within the RecET superfamily. This trimeric assembly contrasts with the canonical C4-symmetric homotetramer of *E. coli* RecE but parallels the architecture of *E. coli* Redα. However, unlike Redα, Che9cRecE and its homologs share greater monomeric fold similarity with RecE, placing them as an evolutionary intermediate between RecE tetramers and Redα trimers.

Several important questions remain for future investigation. First, the moderate resolution of our cryo-EM reconstruction precludes atomic-level analysis of Che9cRecECL dynamics and its precise interactions with DNA. High-resolution structures of Che9cRecE in complex with dsDNA and ssDNA substrates will be essential to rigorously test our electrostatic and steric regulation model. Second, the contribution of Che9cRecT to the high efficiency of Cas-ET remains largely unexplored; future work should examine how Che9cRecT promotes strand exchange and interacts with host replication and repair pathways. Finally, while Cas-ET functions effectively in two model *Streptomyces* strains, its performance across genetically diverse actinobacteria, including industrially important natural product producers, needs comprehensive evaluation.

In conclusion, our work delivers a valuable genome-editing tool for bacteria and uncovers a previously unappreciated layer of regulation in the RecE family. The discovery that C-termini act as tunable rheostats for exonuclease activity — and that these modules can be ported across diverse exonucleases to boost recombination efficiency — provides both a mechanistic framework and a general engineering strategy for advancing recombineering technology.

## Methods

### BLASTp search and protein sequence clustering

BLASTp searches were performed against the PhagesDB ^52^ and NCBI BLAST ^53^ databases using *E. coli* RecE (NCBI accession: WP_000105143.1) as the query sequence. Candidate RecE homologs from actinobacteriophages were retained if they harboured DUF3799 domains as annotated in the InterPro database ^54^. Their cognate RecT homologs were subsequently identified manually via BLASTp searches and InterPro domain annotation.

### Strains and culture conditions

*E. coli* strains were cultured at 37 °C in Luria–Bertani (LB) medium supplemented with appropriate antibiotics: ampicillin, 100 μg ml⁻¹; chloramphenicol, 15 μg ml⁻¹; Zeocin, 15 μg ml⁻¹; or apramycin, 20 μg ml⁻¹. *S. coelicolor* A3(2) and *S. roseosporus* NRRL 15998 were grown at 30 °C on mannitol–soy flour (MS) agar plates supplemented with appropriate antibiotics: hygromycin, 50 μg ml⁻¹; apramycin, 50 μg ml⁻¹; nalidixic acid, 50 μg ml⁻¹. *P. putida* KT2440 was cultivated at 30 °C in LB medium supplemented with appropriate antibiotics: kanamycin, 15 μg ml⁻¹; gentamicin, 30 μg ml⁻¹.

### Construction of plasmid for *Streptomyces* genome editing

The pCas-hyg plasmid was constructed from pCRISPomyces2 ^29^ via the RedEx seamless site-directed mutagenesis method ^11^ by sequentially inserting two DNA fragments: a 20-bp sequence encoding CRISPR-Cas9 guide RNA spacer targeting *actVA5* of *S. coelicolor* A3(2), and the HA-hyg-HA cassette which comprises a hygromycin resistance gene flanked by 1,500-bp homology arms flanking the *actVA5* locus.

The 31 actinobacteriophage RecET recombinase gene pairs were synthesized by Genscript (Suzhou, China) and individually inserted into pCas-hyg via the RedEx seamless site-directed mutagenesis method ^11^ to generate pCas-actRecET-hyg plasmids.

The pCas-hyg-HA plasmids for Fig. 1d was constructed from pCas-hyg via the RedEx seamless site-directed mutagenesis method ^11^ by individually inserting the HA-hyg-HA cassette which comprises a hygromycin resistance gene flanked by 50-, 100-, 200-, 500-, 1,000-, and 1,500-bp homology arms flanking the *actVA5* locus.

The pCas-Che9cRecET-hyg-HA plasmids for Fig. 1d was constructed from pCas-Che9cRecET-hyg via the RedEx seamless site-directed mutagenesis method ^11^ by individually inserting the HA-hyg-HA cassette which comprises a hygromycin resistance gene flanked by 50-, 100-, 200-, 500-, 1,000-, and 1,500-bp homology arms flanking the *actVA5* locus.

pBACspn-CasChe9cRecET-apra-HA for Fig. 1e was constructed from pBACspnNEW ^55^ harboring the 79-kb spinosad gene cluster via linear-circular recombineering mediated Redαβ ^8^ by sequentially inserting two DNA cassettes: the HA-zeo cassette which comprises a zeomycin resistance gene and 50-, 100-, 200-, 500-, 1,000-, and 1,500-bp homology arms upstream of the *actVA5* locus, and the CasChe9cRecET-apra-HA cassette which comprises the *cas9* gene, the *che9cRecET*genes, the 20-bp sequence encoding CRISPR-Cas9 guide RNA spacer targeting *actVA5* of *S. coelicolor* A3(2), the apramycin resistance gene, and 50-, 100-, 200-, 500-, 1,000-, and 1,500-bp homology arms downstream of the *actVA5* locus.

pBACspn-CasChe9cRecET-cda, pBACspn-CasChe9cRecET-che, pBACspn-CasChe9cRecET-phi, pBACspn-CasChe9cRecET-pro, and pBACspn-CasChe9cRecET-myc plasmids for Fig. 1f harbor the 79-kb spinosad gene cluster and target the the calcium-dependent antibiotic gene cluster, the coelichelin gene cluster, the *phiC31-attB* site, the prodigiosin gene cluster, and the coelimycin gene cluster in the chromosome of *S. coelicolor* A3(2), respectively. They were constructed from pBAC-spnNEW via linear-circular recombineering mediated Redαβ ^8^ by sequentially inserting two DNA cassettes: the HA-zeo cassette which comprises a zeomycin resistance gene and the 1,000-bp homology arm upstream of above 5 loci, and the CasChe9cRecET-apra-HA cassette which comprises the *cas9* gene, the *che9cRecET*genes, the 20-bp sequence encoding CRISPR-Cas9 guide RNA spacer targeting above 5 loci, the apramycin resistance gene, and the 1,000-bp homology arm downstream of above 5 loci.

The pCas-hyg-dptA plasmid for Fig. 1g was constructed from pCRISPomyces2 ^29^ via the RedEx seamless site-directed mutagenesis method ^11^ by sequentially inserting two DNA fragments: a 20-bp sequence encoding CRISPR-Cas9 guide RNA spacer targeting the *dptA* gene of *S. roseosporus* NRRL 15998, and the HA-hyg-HA cassette which comprises a hygromycin resistance gene flanked by 50-bp homology arms flanking the *dptA* gene. The pCas-Che9cRecET-hyg-dptA plasmid for Fig. 1g was constructed from pCas-hyg-dptA via the RedEx seamless site-directed mutagenesis method ^11^ by inserting the Che9cRecET genes.

### Intergeneric conjugation of *Streptomyces* strains

Plasmids were first electroporated into *E. coli* ET12567 (pUZ8002) ^56^, and subsequently introduced into *Streptomyces* strains via intergeneric conjugation. Positive genome-editing exconjugants were screened on MS agar plates supplemented with nalidixic acid plus hygromycin or apramycin, and further verified by PCR amplification of the homologous recombination junction regions. Genome-editing efficiency was calculated as the number of colony-forming units per 10^7^ *Streptomyces* spores used for each conjugation.

### Fermentation of *S. coelicolor* A3(2) strains and HPLC-MS analysis of spinosyn A production

*S. coelicolor* A3(2) recombinant strains carrying the spinosyn biosynthetic gene cluster were inoculated into 30 mL of tryptic soy broth (TSB) in 250 mL flasks. Following incubation at 30 °C with shaking at 220 rpm for 2 days, 1 mL of the seed culture was transferred to 30 mL of fermentation medium (4% glucose, 1% glycerol, 3% soluble starch, 1.5% soytone, 1% beef extract, 0.65% peptone, 0.05% yeast extract, 0.1% MgSO_4_, 0.2% NaCl, 0.24% CaCO_3_) in 250 ml flasks. Cultures were subsequently incubated at 30 °C with shaking at 220 rpm for 7 days. Next, 1 mL (2% v/v) of Amberlite XAD-16 resin was added to each culture, and the mixture was incubated for a further 2 days. Cells and XAD-16 resin were collected by centrifugation, and the resulting pellet was extracted with 50 mL of methanol. The extract was concentrated to dryness under reduced pressure and reconstituted in 1 mL of methanol.

HPLC-MS analysis was performed on a Bruker Impact HD micrOTOF Q III mass spectrometer equipped with an electrospray ionization source operated in positive ion mode, with 2 μL of extract injected per run. Mass spectra were acquired in centroid mode across a mass range of 100–1500 m/z. Chromatographic separation was achieved on an Acclaim RSLC 120 C18 column (2.1 × 100 mm) using gradient elution with mobile phase A (H_2_O supplemented with 0.1% formic acid) and mobile phase B (acetonitrile supplemented with 0.1% formic acid) at a constant flow rate of 0.3 mL/min. UV absorbance was monitored across 200–600 nm. The gradient elution program for spinosyn A was set as follows: 0–5 min, linear gradient from 5% to 50% B; 5–20 min, isocratic elution at 50% B; 20–25 min, linear gradient from 50% to 95% B; 25–30 min, isocratic elution at 95% B; 30–35 min, column re-equilibration at 5% B.

### Structure prediction and analysis

Three-dimensional structures of all actinobacteriophage RecE homologs, homotrimeric assemblies of Che9cRecE and BogieRecE, and Che9cRecE–DNA and BogieRecE–DNA trimer complexes were predicted using AlphaFold3 ^38^. Five structural models were generated for each target; owing to high structural concordance and consistent confidence scores across models, the top-ranked AlphaFold3 model was selected for downstream analysis. Structural visualization and analysis were performed using PyMOL v3.1.5.1 (Schrödinger, LLC).

### Structural similarity matrix construction, hierarchical clustering, and heatmap visualization

Pairwise structural alignments of all protein structures were performed using TM-align ^57^. TM-scores are length-normalized and direction-dependent, with values differing depending on which structure is used as the reference ^57, 58^. We therefore computed a symmetric average TM-score (TM_avg_) for each pair of structures (*i*, *j*) as follows:

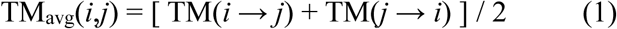

To account for local geometric divergence between aligned residues, each TM_avg_ value was weighted by an exponential RMSD penalty term to yield a pairwise similarity score:

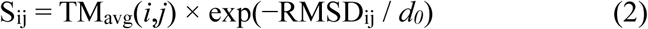

where *d_0_* = 5.0 Å is the empirical distance threshold used by TM-align to define structurally matched residue pairs. Diagonal elements of matrix *S* were set to 1. Off-diagonal elements were min–max normalized to the range [0,1] and subsequently symmetrized to generate the final normalized similarity matrix *S*_norm_. A structural distance matrix *D* was then derived as:

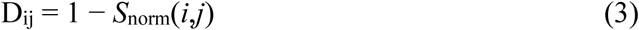

with all diagonal elements set to 0.

Hierarchical clustering was performed on matrix *D* using the unweighted pair-group method with arithmetic mean (UPGMA, average linkage) ^39^, as implemented in SciPy (scipy.cluster.hierarchy.average) ^59^. The resulting dendrogram was exported in Newick format and visualized in FigTree (http://tree.bio.ed.ac.uk/software/figtree/). Clade support was assessed via bootstrap resampling of taxa: taxa were resampled with replacement from the distance matrix for 1,000 replicates, and a UPGMA tree was reconstructed for each replicate. Support values are reported as the percentage of replicates in which each clade of the original tree was recovered. Finally, the *S*_norm_ matrix was visualized as a heatmap using Matplotlib ^60^, with rows and columns ordered to match the leaf order of the UPGMA dendrogram, so that pairwise structural similarity is presented in direct alignment with the clustering topology.

### Multiple sequence alignment

Multiple sequence alignment was performed using Clustal Omega ^61^, and the alignment was rendered as figures using ESPript 3 ^62^. Secondary structure elements derived from the predicted three-dimensional structural models were mapped onto the aligned sequences.

### Protein expression and purification

The genes encoding actinobacteriophage RecE homologs were codon-optimized expression in *E. coli* BL21 (DE3) strain. pET-28a plasmids carrying genes encoding N-terminally 6×His-SUMO-tagged target proteins were electroporated into *E. coli* BL21 (DE3). Transformants were cultured in LB medium supplemented with 50 μg mL⁻¹ kanamycin at 37 °C with shaking until the OD_600_ reached 0.6–1.0. Protein expression was induced with IPTG at a final concentration of 0.2 mM, followed by further incubation at 18 °C for 16 h. Cells were harvested by centrifugation, resuspended in lysis buffer (20 mM Tris-HCl pH 8.0, 300 mM NaCl, 5% glycerol, 10 μg mL^-1^ DNase I, 1 mM PMSF) and disrupted by sonication on ice. Insoluble cell debris was removed by centrifugation at 15,000 × *g* for 90 min at 4 °C. The supernatant was loaded onto a Ni-NTA affinity column (C600793, BBI) pre-equilibrated with lysis buffer. The column was washed with five column volumes of wash buffer (20 mM Tris-HCl pH 8.0, 300 mM NaCl, 5% (v/v) glycerol), and the target protein was subsequently eluted using elution buffer (20 mM Tris-HCl pH 8.0, 300 mM NaCl, 5% (v/v) glycerol, 250 mM imidazole).

The N-terminal 6×His-SUMO tag was cleaved by Ulp1 protease during overnight incubation at 4 °C, and removed via reverse Ni²⁺ affinity chromatography on a HisTrap HP column. The resulting tag-free protein was loaded onto a HiTrap Q FF anion-exchange column (cat. no. 17515601, Cytiva) and eluted with a linear 0–1 M NaCl gradient. Finally, pooled elution fractions were concentrated and further purified by size-exclusion chromatography using a Superdex 200 Increase 10/300 GL column (cat. no. 28990944, Cytiva). Protein purity was assessed by SDS–PAGE, and protein concentration was quantified using a NanoDrop spectrophotometer (cat. no. NP80, IMPLEN).

### Protein electrophoretic mobility shift assays

Complementary oligonucleotides were mixed an equimolar ratio, heated at 95 °C for 10 min and gradually cooled to 25 °C to yield partial-duplex dsDNAs.

dsDNA-1: 5′OH-CGTACCGGTGGCGCACG-3′OH

5’OH-CGTGCGCCACCGGTA-3’OH.
dsDNA-2: 5′PO₄-CGTACCGGTGGCGCACG-3′OH

5’OH-CGTGCGCCACCGGTA-3’OH.
dsDNA-3: 5′OH-AAGTGGGCGGCGAAACC-3′OH

5’OH-GGTTTCGCCGCCCAC-3’OH.
dsDNA-4: 5′PO₄-AAGTGGGCGGCGAAACC-3′OH

5’OH-GGTTTCGCCGCCCAC-3’OH.

A 30 μL reaction mixture containing 4.2 μM Che9cRecE, 4.2 μM dsDNA, 20 mM Tris-HCl (pH 8.0), 150 mM NaCl, 1 mM DTT, 2 mM EDTA, and 10% (v/v) glycerol was incubated at 25 °C for 20 min. The reaction samples were subsequently analyzed by native polyacrylamide gel electrophoresis.

### Exonuclease activity assays in the buffer containing different ions

A 30 μL reaction mixture containing 4.2 μM Che9cRecE, 4.2 μM dsDNA, 20 mM Tris-HCl (pH 8.0), 150 mM NaCl, 1 mM DTT, 10% (v/v) glycerol, and 1 mM MgCl_2_, 1 mM CaCl_2_, or 2 mM EDTA was incubated at 25 °C for 40 min. The reaction samples were subsequently analyzed by native polyacrylamide gel electrophoresis.

### Exonuclease activity assay

The exonuclease assay was adapted from a previously described method ^24^. Reactions were carried out in a 10 μL mixture containing 1.2 nM BamHI-linearized 2,686-bp pUC19 DNA (corresponding to 2.4 nM DNA ends), 67 mM glycine-KOH buffer (pH 9.4), 2.5 mM MgCl_2_, 50 μg mL⁻¹ BSA, and trimeric exonuclease at either 1.6 nM or 0.6 nM. Incubations were performed at 30 °C or 37 °C. Reactions were quenched by adding 1 μL of 250 mM EDTA. Subsequently, 2 μL of BlueJuice™ Gel Loading Buffer (cat. no. 10816015, Invitrogen) was added to each sample, and reaction products were resolved by electrophoresis on a 0.8% agarose gel prestained with SYBR Gold (cat. no. S11494, Invitrogen). Denatured ssDNA, prepared by heating 0.6 nM BamHI-linearized 2,686-bp pUC19 dsDNA at 95 °C, was run alongside as a reference marker to indicate the migration position of the ssDNA digestion product.

### Cryo-EM data collection and processing

Purified Che9cRecE was diluted to 0.96 mg mL⁻¹. A 4 μL aliquot of the protein sample was applied to M01-Au300-1.2/1.3 300-mesh cryo-EM grids (Nanodim Technology Co.), which had been glow-discharged at 15 mA for 60 s. Grids were immediately plunge-frozen in liquid ethane using a Vitrobot Mark IV (Thermo Fisher Scientific) under the following conditions: 4 °C, 100% relative humidity, 3 s blot time, blot force 3, and 10 s wait time.

Cryo-EM micrographs were acquired on a Titan Krios G4 transmission electron microscope operated at 300 kV in super-resolution mode. Automated data collection was performed using EPU software, with a defocus range of −1.6 to −2.6 μm, a nominal magnification of 105,000×, and a calibrated pixel size of 0.85 Å. The dose rate was set to 50 e⁻ Å⁻² s⁻¹, and each movie was recorded as a stack of 32 frames. In total, 4,979 movies were collected.

All image processing was performed using cryoSPARC v4.7.0 ^40^. Following motion correction and contrast transfer function (CTF) estimation, 4,207,765 particles were extracted and subjected to multiple rounds of 2D classification. A total of 2,290,128 particles were retained after the final round of 2D classification. These particles were then subjected to iterative rounds of homogeneous refinement and non-uniform refinement, yielding a final 3D reconstruction with an estimated global resolution of 4.35 Å.

### Construction of pCas-actRecEΔC-Che9cRecECL-RecT and pSC101-BAD-gbaΔC-Che9cRecECL-tet plasmids

Using pCas-PhalmRecET-hyg-HA, pCas-BogieRecET-hyg-HA, pCas-PhegasusRecET-hyg-HA, pCas-ZilizebethRecET-hyg-HA, and pSC101-BAD-gba-tet as backbones, the coding regions corresponding to the residues of PhalmRecE, BogieRecE, PhegasusRecE, and ZilizebethRecE, as well as the residues of Redα indicated by the green line in Fig. 2c, were individually replaced with the sequence encoding Che9cRecECL via the RedEx seamless site-directed mutagenesis method ^11^. This generated four chimeric plasmids, respectively: pCas-PhalmRecEΔC-Che9cRecECL-RecT, pCas-BogieRecEΔC-Che9cRecECL-RecT, pCas-PhegasusRecEΔC-Che9cRecECL-RecT, pCas-ZilizebethRecEΔC-Che9cRecECL-RecT, and pSC101-BAD-gbaΔC-Che9cRecECL-tet.

### Construction of pSC101-BAD-gba-Che9cRecECL-tet and pBBR1-Rha-BA-Che9cRecECL-S-kan

The DNA sequence encoding Che9cRecECL and the vector backbones pSC101-BAD-gba-tet ^7^ or pBBR1-Rha-BAS-kan ^11^ were individually prepared by PCR. Above PCR products were assembled via Gibson assembly and electroporated into *E. coli* GB2005 to generate two chimeric plasmids, pSC101-BAD-gba-Che9cRecECL-tet and pBBR1-Rha-BA-Che9cRecECL-S-kan.

### Construction of pSC101-BAD-gTE-tet and pSC101-BAD-gTE-Che9cRecECL-tet

The vector backbone pSC101-BAD-tet, the *red**γ*** gene, the *recT* gene, and the *recE* gene or the *recE-Che9cRecECL* gene were individually prepared by PCR. Above PCR products were assembled via Gibson assembly and electroporated into *E. coli* GB2005 to generate pSC101-BAD-gTE-tet and pSC101-BAD-gTE-Che9cRecECL-tet.

### Recombineering

Recombineering in *E. coli* GB2005 was performed following previously established protocols ^8^. For linear–circular recombineering, the mixture of 200 ng of a linear PCR fragment carrying a kanamycin resistance gene flanked by 50-bp homology arms and 200 ng of circular p15A-cm plasmid ^6^ were used as the substrate. For linear–linear recombineering, the mixture of 200 ng of a linear PCR fragment carrying a kanamycin resistance gene flanked by 50-bp homology arms and 200 ng of a linear PCR fragment of p15A-cm vector ^6^ were used as the substrate.

Linear–circular recombineering in *P. putida* KT2440 was carried out following a previously described protocol ^12^. Three micrograms of a linear PCR product containing a gentamicin resistance gene flanked by 100-bp homology arms was used as the substrate to replace the BAS operon in pBBR1-Rha-BAS-kan or the BA-Che9cRecECL-S operon in pBBR1-Rha-BA-Che9cRecECL-S-kan.

Positive recombinants were selected on LB agar plates supplemented with corresponding antibiotics and further verified by restriction analysis. Recombineering efficiency was calculated as the number of colony-forming units per 10⁹ competent cells per electroporation reaction.

## Acknowledgements

This work was supported by the National Natural Science Foundation of China (32400022 to J.L., 32670034 to H.W.); Natural Science Foundation of Shandong Province (ZR2022JQ11 to H.W.); the Fundamental Research Funds of Shandong University (2023QNTD001 to H.W.); the National Key Research & Development Program of China (2025YFA0922600 to H.W.); Young Talent Development Program of SKLMT (M2025YA04 to H.W.); the Fund for Distinguished Young Scholars of SDU (H.W.); the SKLMT Frontiers and Challenges Project (SKLMTFCP-2023-05 to H.W.). The authors would like to thank Jingyao Qu, Jing Zhu and Zhifeng Li of the Core Facilities for Life and Environmental Sciences, State Key Laboratory of Microbial Technology of Shandong University for their help and guidance in LC-MS.

## Author Contributions Statement

H.W., J.L. and C.J. conceived and designed the research. H.W., J.L., C.J., R.G., and C.W. designed experiments. C.J., R.G., J.L., C.W., and S.W. performed experiments. H.W., J.L., C.J., R.G., C.W., A.F.S., J.F., and Y.Z. discussed the data. H.W., J.L., C.J., R.G., and C.W. wrote the manuscript.

