## Supplemental Figures and Tables for "A portable C-terminal rheostat calibrates RecE/Redα exonuclease activity for enhanced bacterial recombineering"

**Supplementary Table 1. Protein Accession numbers**

| Accession numbers | Proteins |
|---|---|
| NCBI WP_000105143.1 | *E. coli* RecE |
| NCBI WP_270826003.1 | *E. coli* RecT |
| NCBI NP_040618.1 | *E. coli* Redγ |
| NCBI WP_000186853.1 | *E. coli* Redα |
| UniProtKB P03698 | *E. coli* Redβ |
| NCBI QPX61810.1 | Mycobacterium phage CamsterRecE |
| NCBI QPX61811.1 | Mycobacterium phage CamsterRecT |
| NCBI QYW01346.1 | Mycobacterium phage JoleneRecE |
| NCBI QYW01347.1 | Mycobacterium phage JoleneRecT |
| NCBI YP_009013064.1 | Mycobacterium phage BabsiellaRecE |
| NCBI YP_009013065.1 | Mycobacterium phage BabsiellaRecT |
| NCBI YP_009012296.1 | Mycobacterium phage BigNuzRecE |
| NCBI YP_009012297.1 | Mycobacterium phage BigNuzRecT |
| NCBI NP_817737.1 | Mycobacterium phage Che9cRecE |
| NCBI NP_817738.1 | Mycobacterium phage Che9cRecT |
| NCBI YP_009004417.1 | Mycobacterium phage DonovanRecE |
| NCBI YP_009004418.1 | Mycobacterium phage DonovanRecT |
| NCBI YP_009964848.1 | Mycobacterium phage FirstPlacePfuRecE |
| NCBI YP_009964849.1 | Mycobacterium phage FirstPlacePfuRecT |
| NCBI YP_009964534.1 | Mycobacterium phage GreaseLightninRecE |
| NCBI YP_009964535.1 | Mycobacterium phage GreaseLightninRecT |
| NCBI YP_010088225.1 | Mycobacterium phage HCRecE |
| NCBI YP_010088226.1 | Mycobacterium phage HCRecT |
| NCBI YP_009006648.1 | Mycobacterium phage MichelleMyBellRecE |
| NCBI YP_009006649.1 | Mycobacterium phage MichelleMyBellRecT |
| NCBI YP_009198390.1 | Mycobacterium phage PhayonceRecE |
| NCBI YP_009198391.1 | Mycobacterium phage PhayonceRecT |
| NCBI YP_009965174.1 | Mycobacterium phage PurkyRecE |
| NCBI YP_009965175.1 | Mycobacterium phage PurkyRecT |
| NCBI YP_009124711.1 | Mycobacterium phage SbashRecE |
| NCBI YP_009124712.1 | Mycobacterium phage SbashRecT |
| NCBI YP_009595735.1 | Mycobacterium phage SkinnyPeteRecE |
| NCBI YP_009595736.1 | Mycobacterium phage SkinnyPeteRecT |
| NCBI YP_009965259.1 | Mycobacterium phage XaviaRecE |
| NCBI YP_009965261.1 | Mycobacterium phage XaviaRecT |
| NCBI ASD53676.1 | Mycobacterium phage BogieRecE |
| NCBI ASD53677.1 | Mycobacterium phage BogieRecT |
| NCBI AOT24887.1 | Mycobacterium phage NazoRecE |
| NCBI AOT24888.1 | Mycobacterium phage NazoRecT |
| NCBI QHB41312.1 | Mycobacterium phage PhalmRecE |
| NCBI QHB41313.1 | Mycobacterium phage PhalmRecT |
| NCBI URP21031.1 | Mycobacterium phage PhegasusRecE |

| | |
|---|---|
| NCBI URP21032.1 | Mycobacterium phage PhegasusRecT |
| NCBI QIG57638.1 | Mycobacterium phage StressBallRecE |
| NCBI QIG57639.1 | Mycobacterium phage StressBallRecT |
| NCBI QBI97980.1 | Mycobacterium phage ZilizebethRecE |
| NCBI QBI97981.1 | Mycobacterium phage ZilizebethRecT |
| NCBI ACU41506.1 | Mycobacterium phage HopeRecE |
| NCBI ACU41507.1 | Mycobacterium phage HopeRecT |
| NCBI AEL98514.1 | Mycobacterium phage LiefieRecE |
| NCBI AEL98515.1 | Mycobacterium phage LiefieRecT |
| NCBI WDW19673.1 | Mycobacterium phage LOCV1RecE |
| NCBI WDW19674.1 | Mycobacterium phage LOCV1RecT |
| NCBI AYD84006.1 | Mycobacterium phage GrizzlyRecE |
| NCBI AYD84007.1 | Mycobacterium phage GrizzlyRecT |
| NCBI QFG15050.1 | Mycobacterium phage QueenHazelRecE |
| NCBI QFG15051.1 | Mycobacterium phage QueenHazelRecT |
| NCBI UVT31495.1 | Mycobacterium phage SejanusRecE |
| NCBI UVT31496.1 | Mycobacterium phage SejanusRecT |
| NCBI YP_009017043.1 | Mycobacterium phage CharlieRecE |
| NCBI YP_009017044.1 | Mycobacterium phage CharlieRecT |
| NCBI YP_010104182.1 | Mycobacterium phage XulaRecE |
| NCBI YP_010104183.1 | Mycobacterium phage XulaRecT |
| NCBI AOT24453.1 | Propionibacterium phage DoucetteRecE |
| NCBI AOT24454.1 | Propionibacterium phage DoucetteRecT |
| NCBI AOT24329.1 | Propionibacterium phage B3RecE |
| NCBI AOT24330.1 | Propionibacterium phage B3RecT |
| NCBI YP_009007104.1 | *Pseudomonas aeruginosa* phage Ab31 orf37 |
| NCBI YP_009007105.1 | *P. aeruginosa* phage Ab31 orf38 |
| NCBI YP_009007103.1 | *P. aeruginosa* phage Ab31 orf36 |

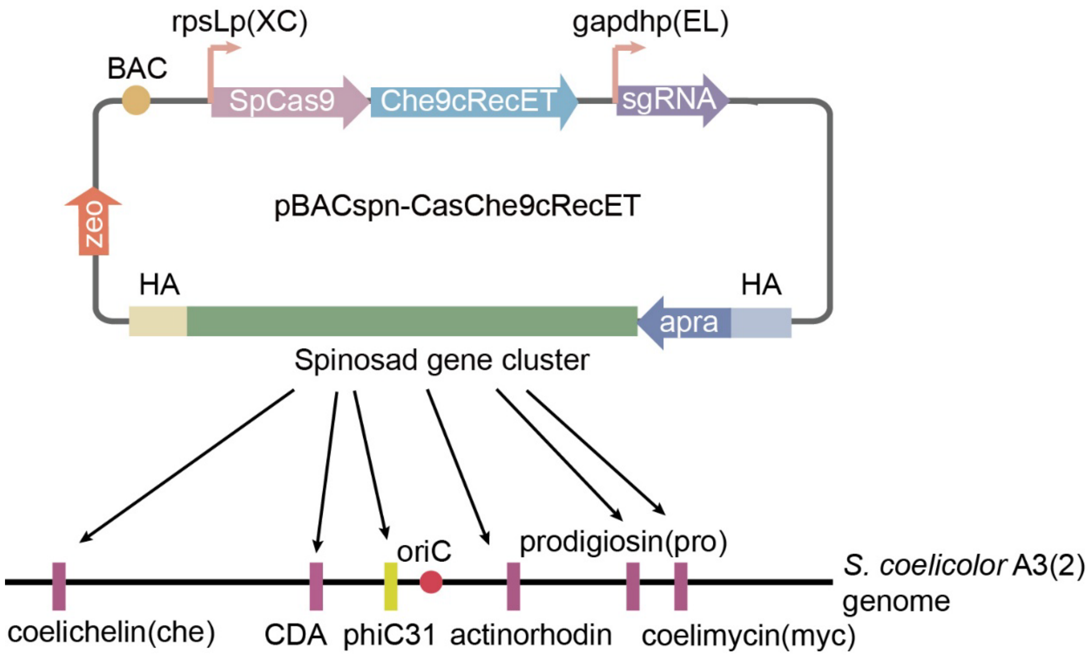


**Supplementary Figure 1. Schematic of insertion of the 79-kb spinosyn gene cluster at the *actVA5* locus of the *S. coelicolor* A3(2) chromosome using Cas-ET$_{Che9c}$.**

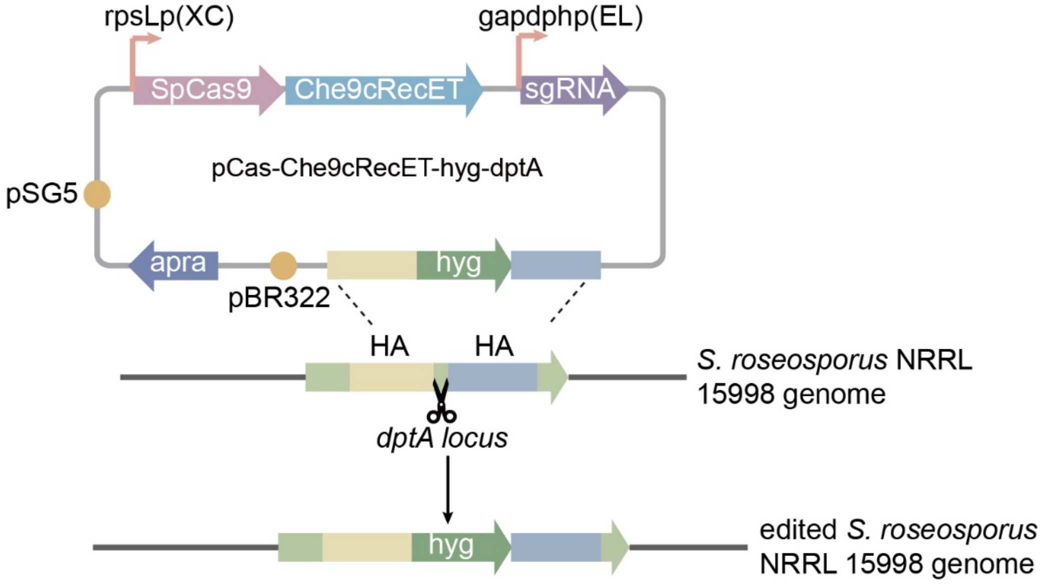


**Supplementary Figure 2. Schematic of insertion of the *hyg* gene into the *dptA* gene of the *S. roseosporus* NRRL 15998 genome using Cas-ET$_{Che9c}$.**

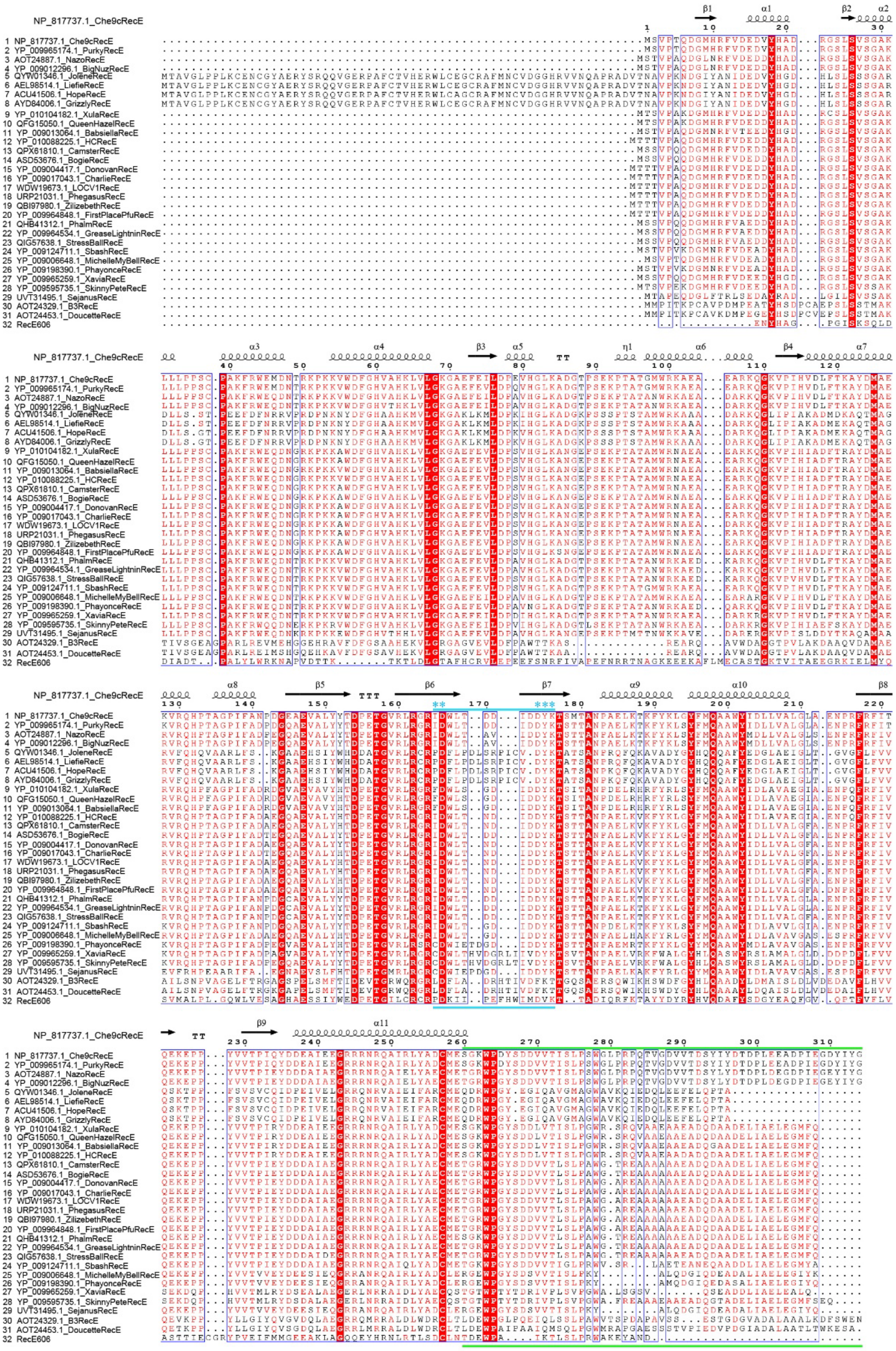


**Supplementary Figure 3. Multiple sequence alignment of full-length amino acid sequences of RecE homologs.** The multiple sequence alignment was performed with the MUSCLE algorithm. Secondary structures of Che9cRecE were indicated above the sequences. The region highlighted in blue was the PD-(D/E)XK domains. The asterisks indicated the active site residues. The region highlighted in green was the Che9cRecE subfamily-specific C-termini.

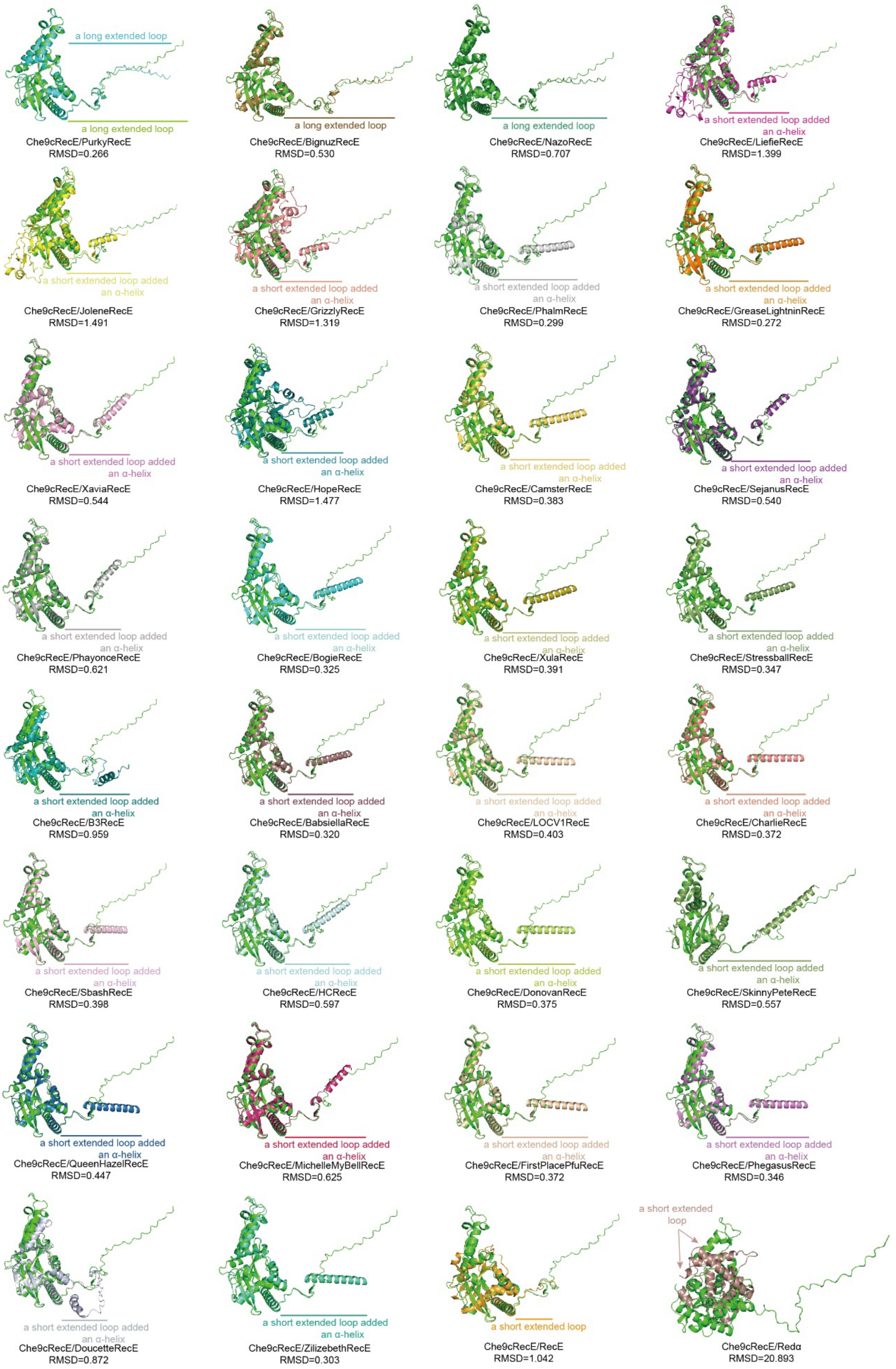


**Supplementary Figure 4. Comparison of protein monomer structures of Che9cRecE and other RecE homologs predicted by AlphaFold3, and *E. coli* RecE (PDB 3H4R) and Redα (PDB 1AVQ).**

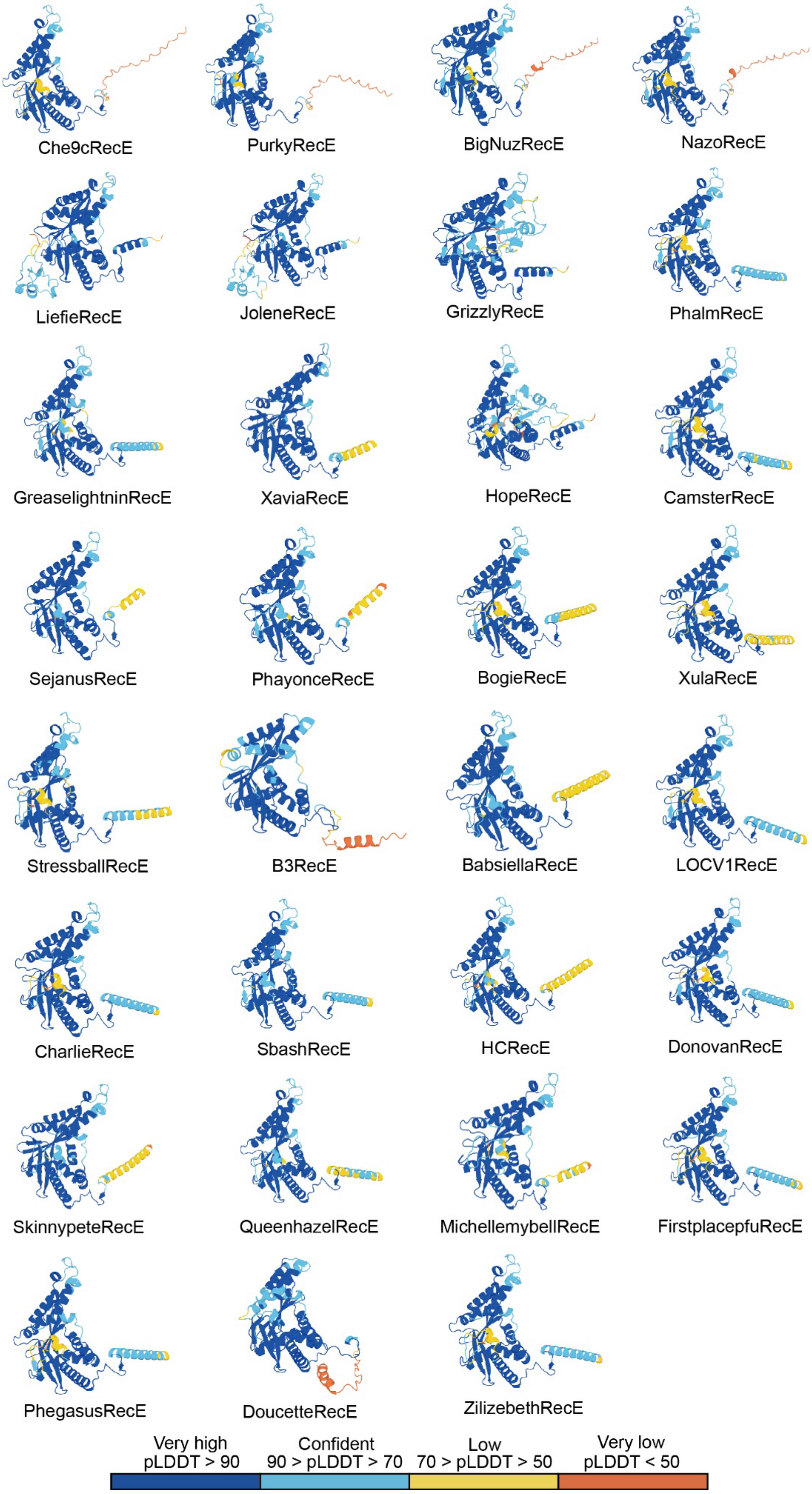


**Supplementary Figure 5. Protein monomer structures of RecE homologs predicted by AlphaFold3 shown with predicted local-distance difference test (pLDDT).**

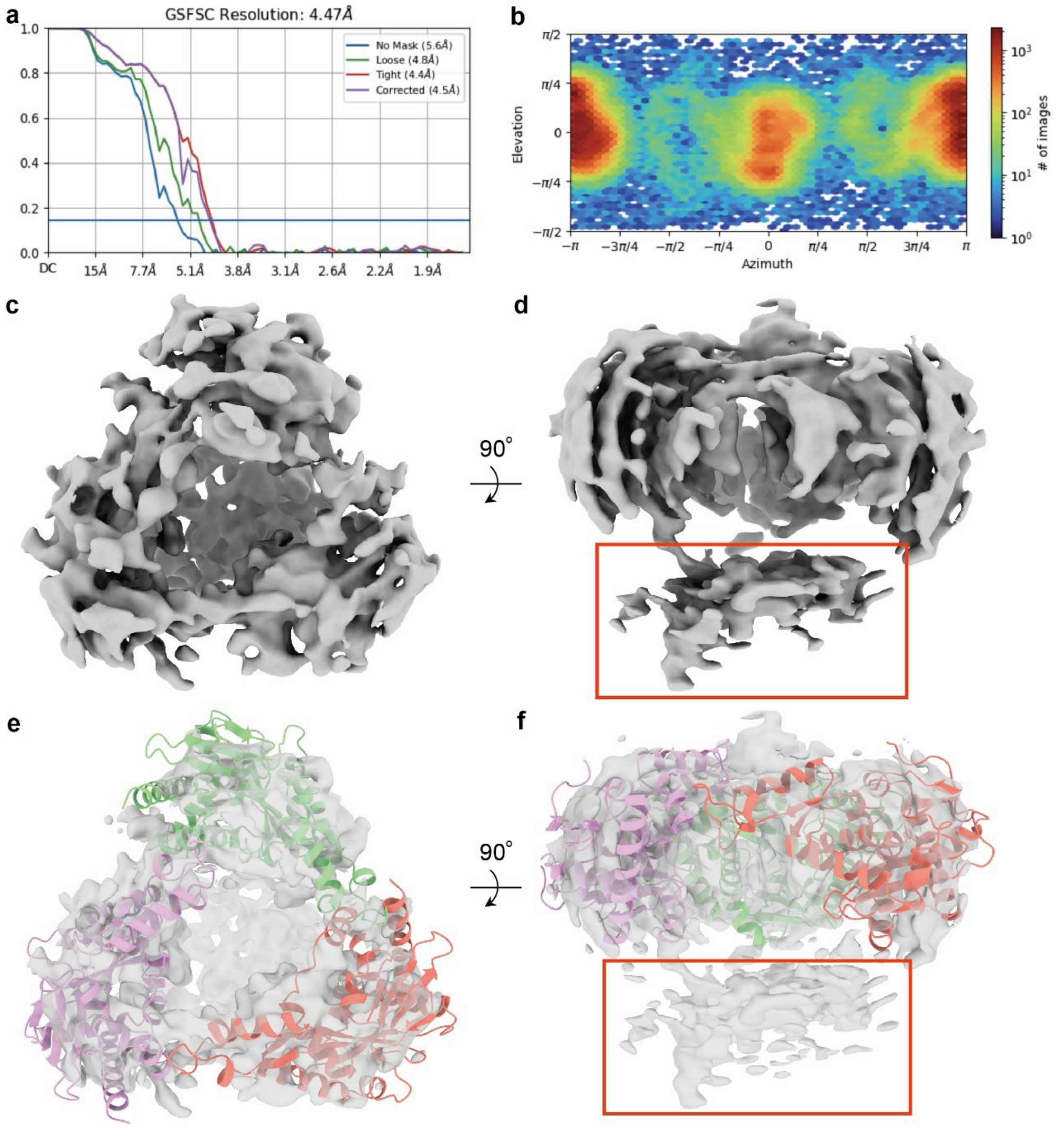


**Supplementary Figure 6. The cryo-EM map of Che9cRecE. a,** Resolution estimate by gold-standard Fourier Shell Correlation (FSC). **b,** Angular distribution of particles used for final reconstruction. **c and d,** Electron density for cryo-EM map of Che9cRecE. Red box indicated the density region hypothesized to be the Che9cRecE C-terminus. **e and f,** Results of fitting the Alphafold3-predicted structure of Che9cRecE into the currently computed 3D map. Red box indicated the density region hypothesized to be the Che9cRecE C-terminus.

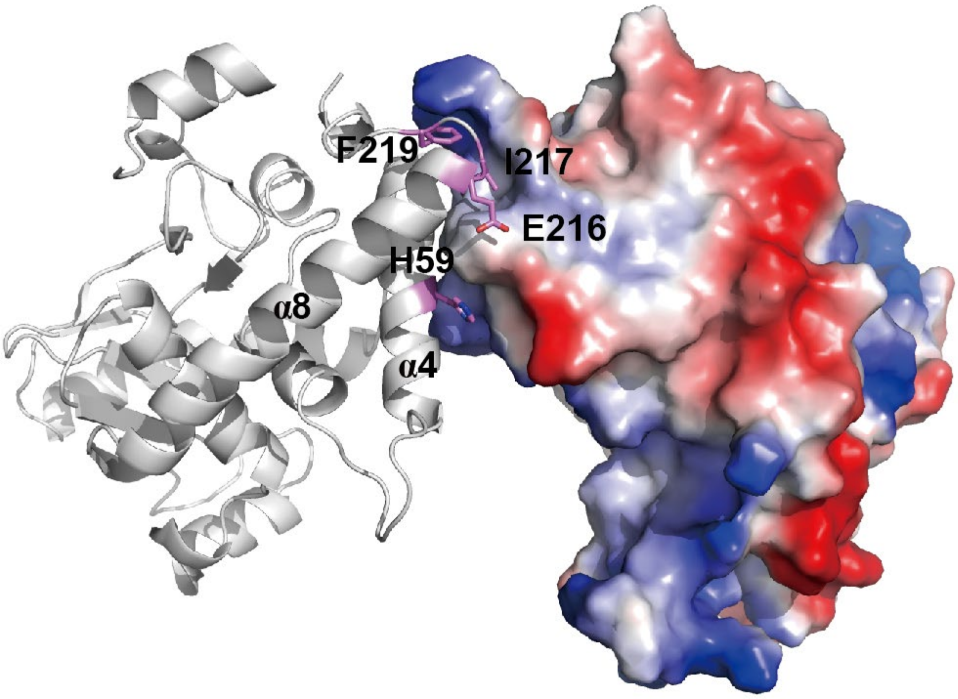


**Supplementary Figure 7. Inter-subunit interface diagrams of *E. coli* Redα.** Hydrophobic core residues at the inter-subunit interfaces were shown in black.

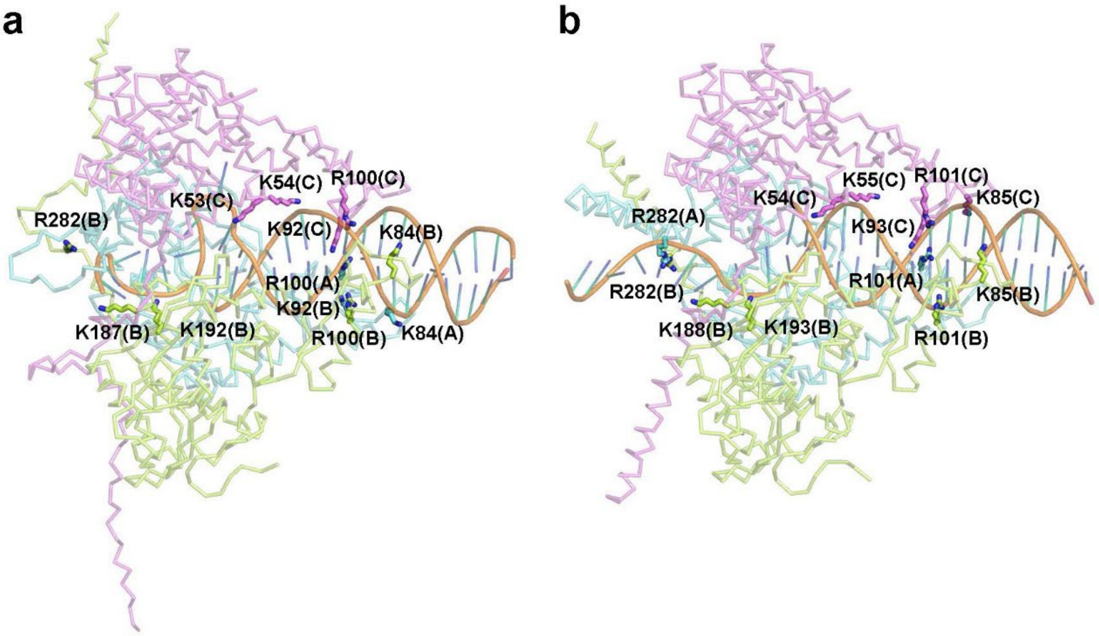


**Supplementary Figure 8. DNA-binding patterns at the residue level of the Che9cRecE–DNA (a) and BogieRecE–DNA (b) complex models.** The views were shown along the central DNA-binding channel. The three subunits of each RecE homotrimer were displayed in different colors (pink, cyan and yellow). A partial dsDNA substrate was represented with an orange backbone and blue base pairs. The side chains of positively charged arginine (Arg) and lysine (Lys) residues that formed salt bridges with the DNA phosphate backbone were shown as sticks and labeled.

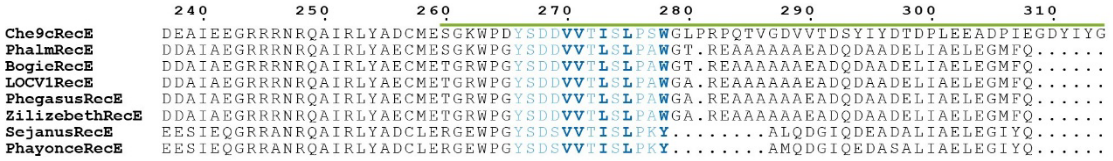


**Supplementary Figure 9. Sequence alignment of the C-terminal regions of RecE homologs**. The C-terminal loop region of Che9cRecE is marked with a green line. Residues located at inter-subunit interfaces are colored blue; hydrophobic core residues are highlighted in bold.

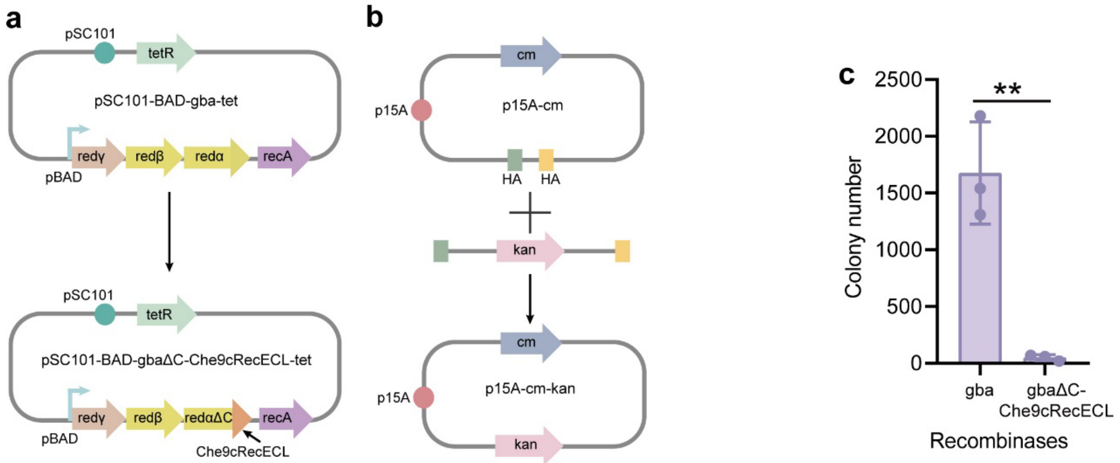


**Supplementary Figure 10. Gene editing efficiency in *E. coli* after replacing the C-terminus of Redα with the Che9cRecECL in Redαβγ system. a,** Schematic diagrams of plasmids plasmids pSC101-BAD-gba-tet and pSC101-BAD-gbaΔC-Che9cRecECL-tet. **b,** Schematic diagram of linear-circular recombineering. **c,** Linear-circular recombineering efficiency mediated by pSC101-BAD-gba-tet and pSC101-BAD-gbaΔC-Che9cRecECL-tet in *E. coli* GB2005.

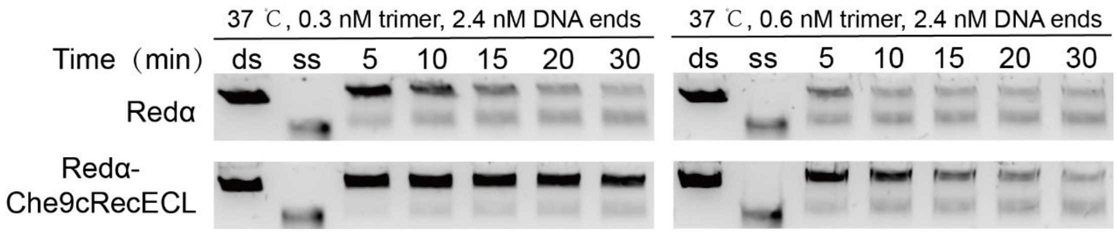


**Supplementary Figure 11. Exonuclease activity assay of Redα and Redα-Che9cRecECL.** Aliquots collected at indicated time post-reaction were analyzed. Lanes ds and ss correspond to the dsDNA substrate and ssDNA product, respectively.
